# ZapA employs a two-pronged mechanism to facilitate Z ring formation in *Escherichia coli*

**DOI:** 10.1101/2024.12.31.629942

**Authors:** Yuanyuan Cui, Han Gong, Di Yan, Hao Li, Wenjie Yang, Ying Li, Xiangdong Chen, Joe Lutkenhaus, Sheng-You Huang, Xinxing Yang, Shishen Du

## Abstract

The tubulin-like protein FtsZ assembles into the Z ring which leads to the assembly and activation of the division machinery in most bacteria. ZapA, a widely conserved protein that interacts with FtsZ, plays a pivotal role in organizing FtsZ filaments into a coherent Z ring. Previous studies revealed that ZapA forms a dumbbell-like tetramer that binds cooperatively to FtsZ filaments and aligns them in parallel, leading to the straightening and organization of FtsZ filament bundles. However, how ZapA interacts with FtsZ remains obscure. In this study, we uncover how ZapA interacts with FtsZ to facilitate Z ring formation in *Escherichia coli*. We find that mutations affecting surface exposed residues at the junction between adjacent FtsZ subunits in a filament as well as in an N-terminal motif of FtsZ weaken its interaction with ZapA *in vivo* and *in vitro*, indicating that ZapA binds to these regions of FtsZ. Consistent with this, ZapA prefers FtsZ polymers over monomeric FtsZ molecules and site-specific crosslinking confirmed that the dimer head domain of ZapA is in contact with the junction of FtsZ subunits. As a result, disruption of the putative interaction interfaces between FtsZ and ZapA abolishes the midcell localization of ZapA. Taken together, our results suggest that ZapA tetramers grab the N-terminal tails of FtsZ and then bind to the junctions between FtsZ subunits in the filament to straighten and crosslink parallel FtsZ filaments into the Z ring.

**Significance:** ZapA is a widely conserved FtsZ-associated protein that promotes the organization of the Z ring, the key cytoskeletal element in the bacterial divisome. Although ZapA is known to crosslink FtsZ filaments, how it interacts with FtsZ remains enigmatic. In this study, we find that *E. coli* ZapA utilizes a dual binding mode in which it binds the junction between FtsZ subunits in a filament and to an N-terminal motif in FtsZ so that it can straighten and crosslink parallel FtsZ filaments simultaneously. Since the junction is formed when FtsZ polymerizes and falls apart when FtsZ depolymerizes, this interaction mode indicates that ZapA employs the polymerization dynamics of FtsZ to organize the filaments into the Z ring.

## Introduction

Cell division is one of the most fundamental processes of life and the unique features of the process in bacteria make it a promising target for the development of novel antibiotics. It starts when polymers of the bacterial tubulin FtsZ coalesce into the Z ring at the future division site, which not only functions as a scaffold for the assembly of the divisome complex but also as a guide for septal peptidoglycan synthesis (sPG) (1–5). Time-lapse imaging of FtsZ fluorescent protein fusions in model organisms such as *Escherichia coli* and *Bacillus subtilis* show that FtsZ filaments first form a diffuse structure at midcell, and then coalesce into a condensed ring with a defined width of about 80-100 nm and a thickness of about 40-50 nm (6–8). Interestingly, super resolution microscopic analysis of the Z ring revealed that it is a highly dynamic discontinuous ring-like structure consisting of clusters of FtsZ filaments anchored to the membrane (9–11). Moreover, FtsZ filaments undergo treadmilling, a directional motion in which subunits add at one end and are released from the opposite end, driven by GTP-dependent polymerization and GTPase-dependent depolymerization of FtsZ, respectively (2, 3). Recent studies found that the treadmilling dynamics of FtsZ filaments is important for them to condense into a mature Z ring and distribute sPG synthetic complexes, consisting of FtsQLBWI, around the division site to construct a smooth septum (7, 8, 12, 13). However, how FtsZ filaments are organized within the Z ring and how their dynamics are coupled with sPG synthesis remain to be elucidated.

In *in vitro* reconstitution systems, FtsZ filaments tethered to the membrane, either by its natural membrane anchors or by adding a membrane targeting sequence, can self-organize into ring-like structures on supported membranes or in liposomes (14–17). However, these reconstituted Z rings are not as condensed and organized as the Z rings *in vivo*, suggesting that additional factors are necessary for the formation of functional Z rings inside cells. Indeed, it is well demonstrated that FtsZ binding proteins (ZBPs), or FtsZ-associated proteins (Zaps), which can crosslink FtsZ filaments into organized structures *in vitro*, play a critical role in Z ring maturation *in vivo* (7, 18, 19). In *E. coli*, five Zap proteins (ZapA, ZapB, ZapC, ZapD and ZapE) have been identified. Each of these proteins, except for ZapB which affects Z ring formation indirectly via ZapA, can interact with FtsZ directly and crosslink FtsZ filaments into large bundles *in vitro* (18, 20–27). The absence of ZapA or ZapB causes abnormal septa and a slight delay in cell division, but the absence of multiple Zap proteins results in a deficiency in forming condensed Z rings, leading to a severe division defect (18, 19). On the other hand, overexpression of any one of these Zap proteins prevents normal Z ring formation in *E. coli*, suggesting that excessive cross-linking of FtsZ filaments is detrimental for proper Z ring organization. In *B. subtilis*, the absence of ZBPs also results in severe defects in Z ring condensation, ultimately leading to divisome assembly and sPG synthesis defects (7). The absence of Zaps or ZBPs in other bacterial species, such as *Caulobacter crescentus* and *Streptococcus pneumoniae*, has also been reported to cause aberrant Z ring formation (28, 29), suggesting that employment of FtsZ crosslinkers to facilitate Z ring organization is widespread in diverse bacterial species.

ZapA is a very widely conserved FtsZ-associated protein within the bacterial kingdom. It was first discovered in *B. subtilis* as a suppressor of MinD overexpression, which prevents Z ring formation and causes lethality (20). Although ZapA is not essential for division, it is synthetic lethal with EzrA or DivIVA in *B. subtilis*, which are positive regulators of Z ring formation, and synthetic sick with other *zap* genes in *E. coli* (18–20, 30, 31). Importantly, ZapA turns out to be a part of the Ter linkage that promotes and stabilizes the Z ring as well as modulates constriction dynamics in *E. coli* (32–34). The Ter signal is a multilayer protein network where ZapB links ZapA bound to FtsZ to MatP which is bound to the terminus region of the chromosome and organizes it into a macrodomain (32, 35). ZapA and ZapB form a cloud-like structure at midcell independent of FtsZ by linkage to MatP and promote rapid formation of the Z ring (36).

DapE, a critical enzyme for the synthesis of the peptidoglycan precursor diaminopimelic acids (DAP), strengthens the Ter signal in a ZapB dependent fashion (37). In *C. crescentus*, the Ter signal consists of ZapA, ZauP and ZapT, the latter two are functional homologs of ZapB and MatP, respectively (28, 38, 39). In the absence of ZapA, ZapB or MatP, FtsZ filaments tend to form loose spiral ring-like structures instead of condensed Z rings, resulting in delayed division and twisted septa (19, 21), suggesting an important role for ZapA (the Ter signal) in organizing the Z ring.

ZapA forms a dumbbell-like tetramer with a dimer head on each end, which can bridge FtsZ filaments to form ladder-like structures and bundles *in vitro* (40, 41). Tetramerization is essential for ZapA function as mutations disrupting tetramer formation inactivate ZapA (42, 43). Several studies have found that ZapA interacts with FtsZ in a 1:1 ratio and mutations in the dimer head of ZapA reduce its interaction with FtsZ (40, 41, 44–46), suggesting that the dimer head is the binding site for FtsZ. However, where and how a ZapA tetramer contacts FtsZ is still mysterious. If each subunit of the ZapA tetramer binds to an FtsZ molecule, the two ZapA monomers in a dimer head need to bind distinct sites on two adjacent FtsZ molecules in a filament because of the 2-fold symmetry of the ZapA dimer. Also, given the symmetry of the bipolar ZapA tetramer, the orientation of the two FtsZ filaments have to be anti-parallel (40). However, this is in conflict with the directional motion of FtsZ filament patches in mature Z rings as observed by advanced light microscopy and simulation studies (2, 7, 47). A recent *in vitro* reconstitution study of FtsZ filament networks with purified FtsZ, FtsA and ZapA on supported membranes showed that ZapA tetramers align FtsZ filaments in a parallel manner, straightens FtsZ filament bundles and increases the spatial order of the filament networks (46). Moreover, ZapA binds to FtsZ filaments transiently and does not affect FtsZ filament length or treadmilling speed. These observations appear to be in line with the treadmilling dynamics of FtsZ filament patches within the Z ring *in vivo*. However, it is difficult to postulate how ZapA can align FtsZ filaments in parallel without affecting treadmilling dynamics given the lack of binding site information on FtsZ and the symmetry of the ZapA tetramer.

In this study, we investigated the interaction between FtsZ and ZapA by characterizing FtsZ mutants resistant to ZapA overexpression toxicity. Our results show that these mutations specifically weaken the interaction between FtsZ and ZapA *in vivo* and *in vitro*. Most of the mutated residues are located at the exposed part of the junction between FtsZ subunits in the filament, suggesting that it is likely the binding site for ZapA which only forms when FtsZ polymerizes into a filament and is lost as FtsZ depolymerizes. Consistent with this, we found that ZapA binds to stable FtsZ polymers, which is reduced by the resistant mutations. In addition, we found that ZapA also binds to a second site, an N-terminal motif in FtsZ, and mutations in this motif also disrupt its interaction with ZapA. Taken together, these results suggest a two-pronged mechanism in which ZapA tetramers bind to the junctions between FtsZ subunits and the N-terminal motifs of FtsZ to simultaneously straighten and crosslink FtsZ filaments that are rotated 180° relative to each other in the Z-axis. This model reconciles the contradictory observations about ZapA and sheds new light on the organization of FtsZ filaments within the Z ring.

## Results

### ZapA overexpression prevents Z ring condensation but not the treadmilling dynamics of individual filaments

Overexpression of ZapA is known to result in the formation of aberrant FtsZ structures which ultimately leads to a division block in *E. coli* (48), but how is not clear. To investigate this problem, we examined the localization dynamics of FtsZ upon ZapA overproduction in *E. coli*. To do this, FtsZ-mNeonGreen (FtsZ-mNG) was ectopically expressed from a plasmid from a anhydrotetracycline-inducible promoter to monitor Z rings and ZapA was expressed from a second plasmid from an IPTG-inducible promoter (P_tac_::*zapA*) that provides sufficient ZapA to completely block colony formation in the presence of IPTG (Fig. 1A). As expected, FtsZ-mNG formed condensed rotating Z rings in cells in the absence of IPTG, but formed widened and distorted ring-like structures at presumptive division sites within filamentous cells following induction of ZapA (Fig. 1B-C, and Video S1). Interestingly, the treadmilling velocity of FtsZ filaments was not affected, however, instead of treadmilling perpendicular to the long axis of the cells, we found that the filaments treadmilled in various directions (Fig. 1, D-F). These observations indicate that too much ZapA alters the organization of FtsZ filaments and thus inhibits Z ring condensation, consistent with a recent report that ZapA changes the spatial order of the FtsZ filament network but does not affect FtsZ GTPase activity or filament length *in vitro* (46).

**Fig. 1.**
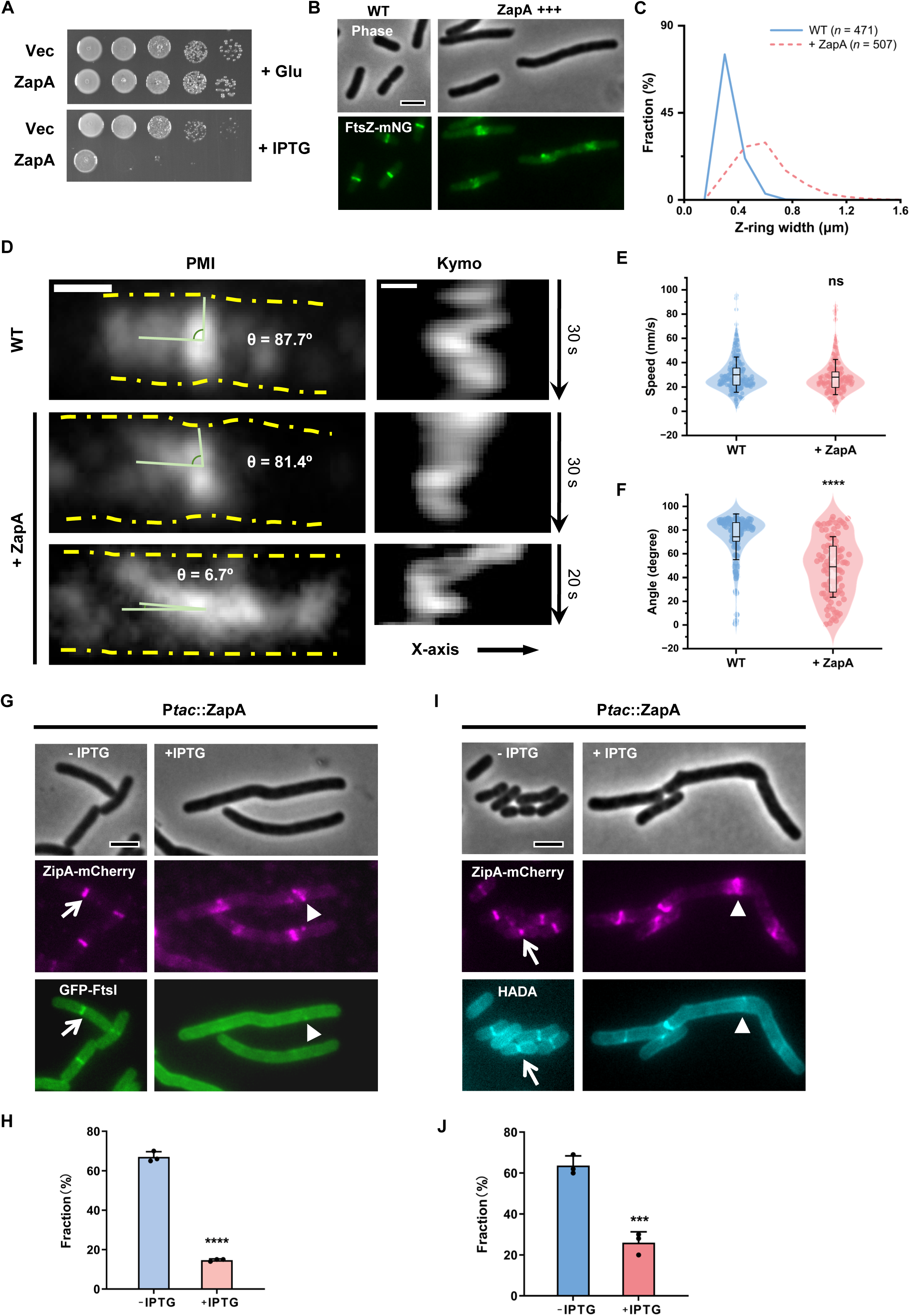
Overexpression of ZapA disrupts Z ring organization and inhibits cell division. (A) Overexpression of ZapA blocks colony formation. Plasmid pEXT22 or pSD319 (*P_tac_::zapA*) was transformed into strain W3110, and ZapA toxicity was assessed with a spot test. A 2 μL aliquot from each 10-fold serial dilution was spotted onto LB plates with Glucose or IPTG and incubated at 37°C overnight before imaging. (B) Representative images of FtsZ-mNG localization in the absence or presence of ZapA overexpression. Strain W3110 was transformed with a plasmid harboring *ftsZ-mNG* under an anhydrotetracycline-inducible promoter and an empty vector or a plasmid carrying *zapA* under the control of an IPTG-inducible promoter. FtsZ-mNG was expressed at the basal level (without ATc) and ZapA was induced with 500 μM IPTG. Samples were taken and immobilized on an LB argarose pad for imaging. (C) Z ring width in the presence or absence of ZapA overexpression. Samples in (B) were analyzed as described in Materials and Methods. Number of Z rings analyzed (WT: n=471, +ZapA: n=507). (D-F) Representative kymographs of FtsZ-mNG (D) and computed FtsZ treadmilling velocity (E) and angles (F) in the absence or presence of ZapA overexpression. Number of filaments analyzed in (E) (WT: n=203, +ZapA: n=193); Number of angles analyzed in (F) (WT: n=94, +ZapA: n=97). (G-H) Representative images of ZipA-mCherry and GFP-FtsI localization (G) and quantitation of their co-localization (H) in the absence or presence of ZapA overexpression. Plasmid pCY129 (*P_tet_::gfp-ftsI*) was transformed into strain CYa35 (W3110, *ftsZ^0^ zipA-mCherry /pACYC, ftsZ & pEXT22,* P_tac_::*zapA*). Cells were grown with or without 500 μM IPTG for 1 h and samples visualized by fluorescence microscopy to assess protein localization. Number of cells analyzed (-IPTG: n=674, +IPTG: n=171). (I-J) Representative images of ZipA-mCherry and HADA localization (I) and quantitation (J) of their co-localization in the absence or presence of ZapA overexpression. The cell cultures of CYa35 (W3110, *ftsZ^0^ zipA-mCherry /pACYC, ftsZ & pEXT22,* P_tac_::*zapA*) were grown with or without 500 μM IPTG for 1 h, and incubated with HADA to label nascent PG. Number of cells analyzed (-IPTG: n=515, +IPTG: n=230). Asterisks denote a significant difference based on a P value of <0.0001 (****) in (F) and (H), and a P value of 0.0008 (***) in (J); ns, not significant. Scale bars, 5 μm.

Since Z ring condensation is critical for efficient recruitment of downstream division proteins to form the complete divisome and for sPG synthesis (7, 8, 19), we checked how ZapA overexpression affected these processes. To do this, we examined the co-localization of ZipA and FtsI, a proxy for the Z ring and a late division protein, respectively, and the co-localization of ZipA and HADA, a fluorescent D-amino acid for tracking nascent PG. We constructed a strain in which *zipA* was replaced with a functional *zipA-mCherry* at its chromosomal locus and GFP-FtsI was ectopically expressed from a plasmid from a anhydrotetracycline-inducible promoter. As shown in Fig. 1G-H, the co-localization of GFP-FtsI and ZipA-mCherry rings was evident in the absence of IPTG, however, the GFP-FtsI signal was greatly reduced at the distorted ZipA-mCherry rings upon induction of ZapA. Moreover, these aberrant structures appeared to be unable to support sPG synthesis efficiently as indicated by the markedly reduced co-localization of HADA with ZipA-mCherry (Fig. 1I-J). Thus, too much ZapA, just like the lack of ZapA, is detrimental for Z ring condensation, ultimately leading to defects in divisome assembly and sPG synthesis.

### Isolation of FtsZ mutants resistant to ZapA overexpression toxicity

To investigate how ZapA interacts with FtsZ, we isolated FtsZ mutants resistant to ZapA overexpression toxicity. This is possible since ZapA is necessary for efficient division but is not essential. We first constructed an FtsZ mutant library by replacing the wild type *ftsZ* in the plasmid pBANG112 with PCR random-mutagenized *ftsZ* as previously described (49, 50). The library was then introduced into the FtsZ depletion strain S17/pKD3C (W3110, ftsZ^0^ / pSC101^Ts^, *ftsZ*), which also harbored the plasmid pSD319 (pEXT22, P_tac_::*zapA*) for ZapA overproduction. Transformants were selected at 42 °C in the presence of 30 μM IPTG, conditions where the pKD3C plasmid is lost and ZapA is overexpressed. Only transformants expressing FtsZ variants that are functional and resistant to overexpressed ZapA could survive (SI Appendix, Fig. S1A).

Using the above approach, we isolated 15 transformants resistant to ZapA overexpression (SI Appendix, Fig. S1B). Sequencing *ftsZ* from the 15 pBANG112^M^ plasmids revealed that they all carried mutations in *ftsZ*, including 9 single mutations, 5 double mutations and a triple mutation (SI Appendix, Table S1). Subsequent analysis of the double and triple mutants revealed that the resistance was mainly due to just one of the substitutions in these mutants (SI Appendix, Fig. S2A). As a result, 13 mutations were isolated by this selection. Mapping these mutations onto a monomeric structure of *E. coli* FtsZ (PDB#: 6UNX) (51) revealed that the vast majority were located on the surface of the polymerization domain, however, two mutations, V128I and V193M, alter residues buried inside the FtsZ structure (SI Appendix, Fig. S2B). Moreover, the surface exposed mutations clustered at the top (the GTP binding pocket) or the bottom part of the FtsZ molecule (containing the T7 loop). A ZapA dimer head is unlikely to contact two sites so far apart in the molecule if it binds to an FtsZ monomer. Intriguingly, when these mutations were mapped onto the filament structure of FtsZ from *Klebsiella pneumoniae* (52), they clustered together at the junction of two adjacent subunits. Three mutations (V128I, V193M and L248M) are a bit farther away but likely affect the conformation of FtsZ (Fig. 2A). This indicates that ZapA may bind to the junction between FtsZ subunits within a filament.

**Fig. 2.**
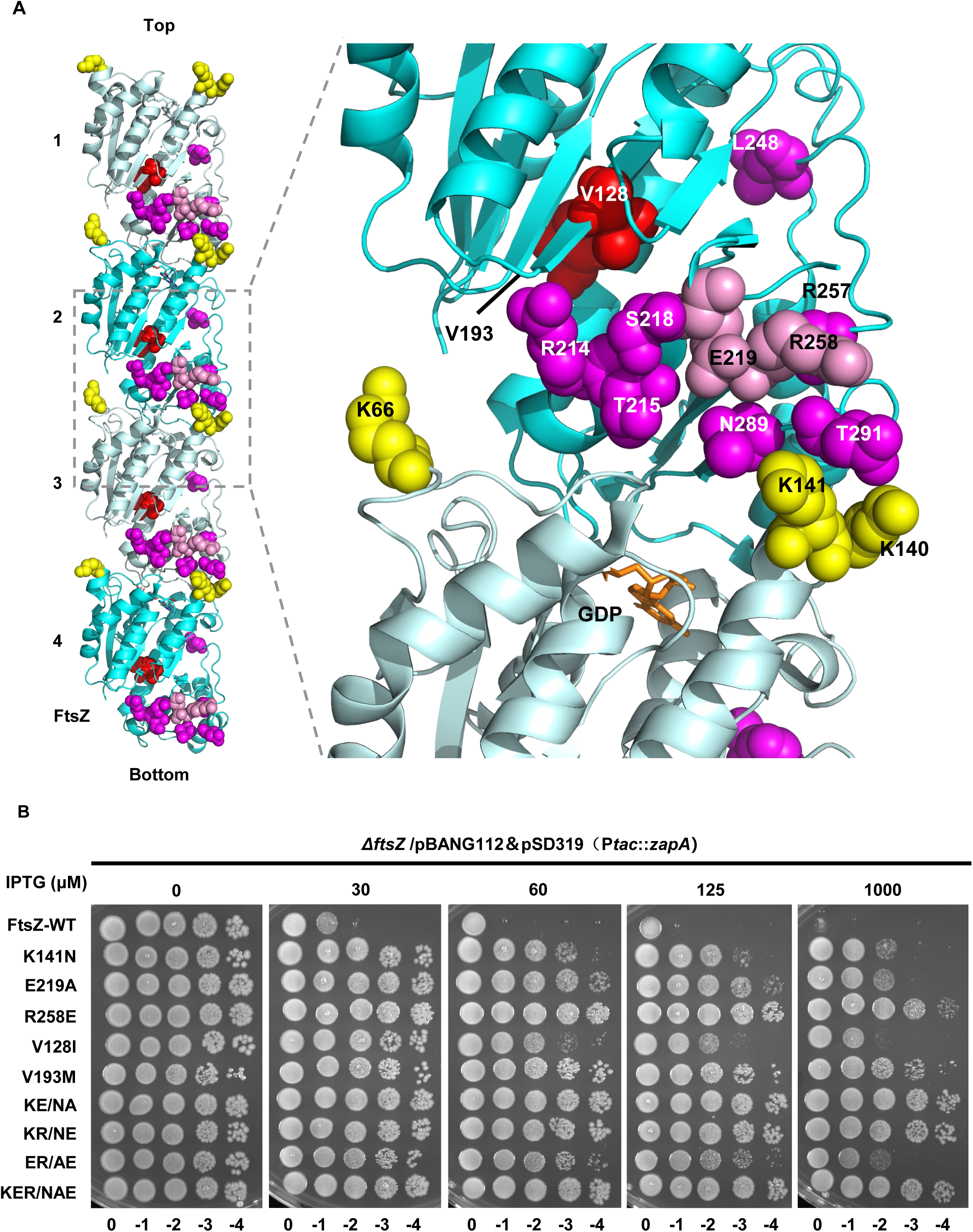
Mutations at the junction of FtsZ subunits in the filament confer resistance to ZapA overexpression. (A) Location of FtsZ mutations in the filament structure of *Klebsiella pneumoniae* FtsZ (PDB# 8IBN). Residues on the surface of FtsZ are colored yellow (top face) or magenta (bottom face), whereas residues buried inside the FtsZ molecule are colored red. Note that mutations isolated by site-directed mutagenesis are colored pink. GMPCPP is shown as stick in brown. Residue numbers are according to *E. coli* FtsZ. (B) Spot test to assess the resistance of FtsZ mutants to ZapA overexpression. Plasmid expressing ZapA (pSD319) was transformed into strains expressing different FtsZ mutants from pBANG112 or its derivatives, and the transformants were subjected to a spot test on plates with or without IPTG.

To test if additional mutations at the junction between FtsZ subunits conferred resistance to ZapA overexpression, we used site-directed mutagenesis to mutate additional residues in this region (SI Appendix, Fig. S3A). Two more resistant mutations (E219A and R258E, both at the bottom part of FtsZ molecule) were isolated, suggesting that the junction between FtsZ subunits indeed contains residues important for interaction with ZapA. Overall, mutations affecting 14 residues were found to provide resistance to ZapA overexpression, with substitutions at 9 providing strong resistance while substitutions at 5 provided modest resistance (SI Appendix, Table S2). Since K141N, E219A, and R258E provide strong resistance to ZapA overexpression and represent mutations on the top and bottom part of the FtsZ molecule that constitute the putative interaction interface, they were chosen for further study (Fig. 2A). V128I, which likely represents a different class of resistant mutation, since it is buried inside of the structure, was also included for the analysis. Reintroduction of the above single mutations or a combination of mutations (KE/NA: K141N, E219A; KR/NE: K141N, R258E; ER/AE: E219A, R258E; KER/NAE: K141N, R258E K141N, E219A, R258E) into pBANG112 revealed that they did not affect the ability of FtsZ to complement the depletion strain S17/pKD3C at 42°C and confirmed they conferred strong resistance to ZapA (Fig. 2B and SI Appendix, Fig. S3). Additionally, cells expressing these FtsZ mutants were still sensitive to the overexpression of ZapC (SI Appendix, Fig. S4), a different Zap protein which also interacts with FtsZ, indicating that they are specifically resistant to ZapA.

### Z rings formed by FtsZ mutants are resistant to the action of ZapA

To explore the resistance phenotype of the FtsZ mutants to ZapA overexpression, we checked if the Z rings formed by these mutants could resist the disrupting effect of ZapA overexpression. To do this, *zipA-mCherry* was introduced into the chromosome of strains expressing wild type FtsZ or the mutants using the λ-Red recombineering system (53). In the absence of IPTG to induce ZapA expression, ZipA-mCherry localized at the division sites in condensed rings (Fig. 3A). Upon overexpression of ZapA, cells expressing wild type FtsZ became filamentous with ZipA-mCherry forming disorganized spiral structures as expected. However, cells expressing FtsZ variants were much shorter and condensed ZipA-mCherry rings were present in most cells, especially in the triple mutant (Fig. 3A and SI Appendix, Fig. S5A). These results demonstrate that resistance to ZapA overexpression is due to its reduced ability to disrupt Z rings formed by these FtsZ mutants. Moreover, quantification of the co-localization of ZipA-mCherry and HADA signal revealed that ZapA overexpresion decreased it from 63% to 26% in cells with wild type FtsZ, but it was not significantly reduced in cells expressing the FtsZ mutants (Fig. 3B-C, SI Appendix, Fig. S5B). Therefore, these FtsZ mutants are able to form functional Z rings that are resistant to overexpressed ZapA, allowing normal divisome assembly and sPG synthesis.

**Fig. 3.**
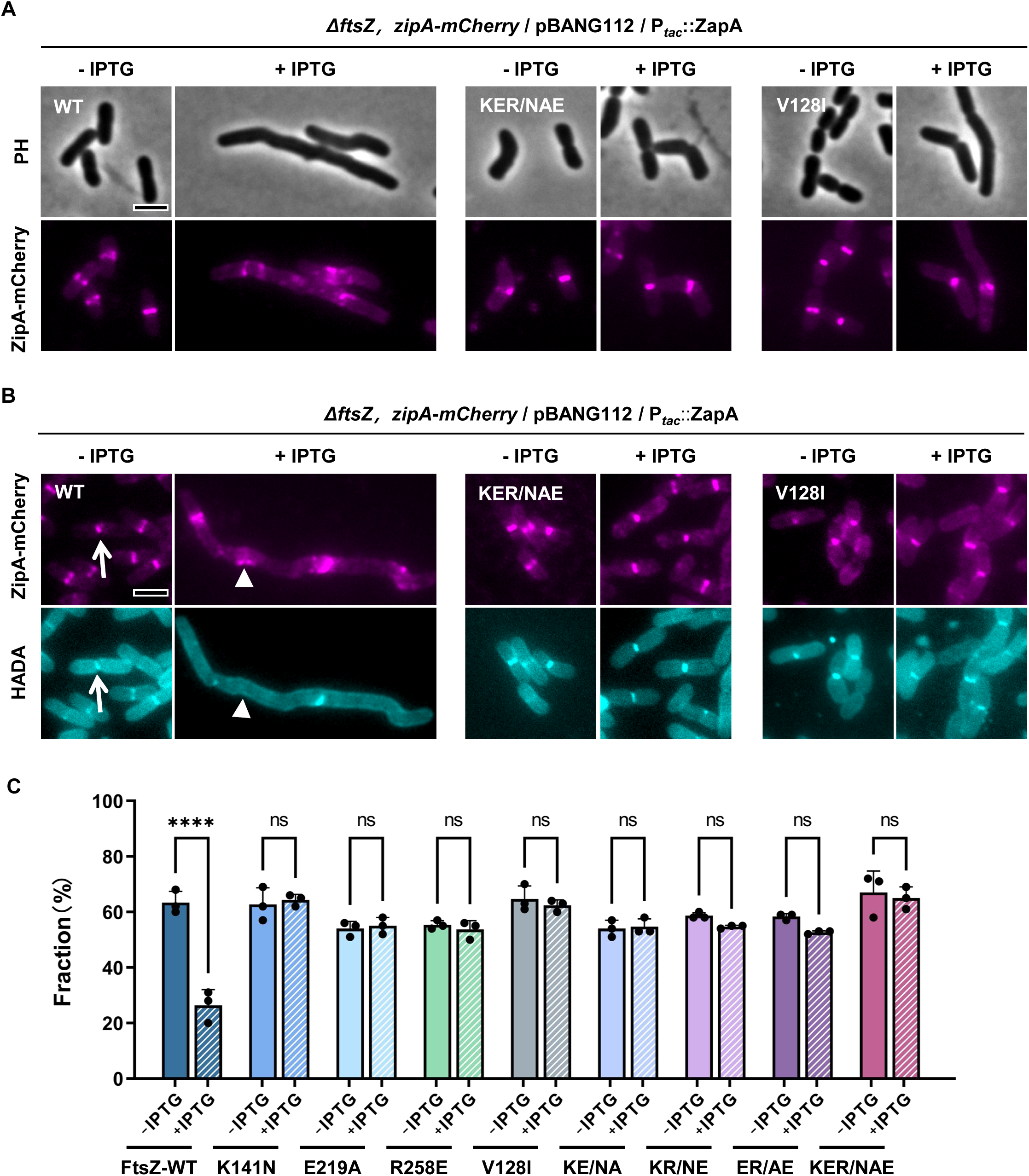
Z rings formed by FtsZ mutants are resistant to ZapA overexpression. (A) Representative images of Z rings (ZipA-mCherry) in cells expressing wild type FtsZ or its variants in the absence or presence of ZapA overexpression. Cells expressing wild type FtsZ or its variants were grown in exponential phase, ZapA was induced with 500 μM IPTG, ZipA-mCherry was expressed from its native promoter. ZipA-mCherry was imaged by fluorescence microscopy. (B) Representative images of co-localization of ZipA-mCherry with HADA in cells expressing wild type FtsZ or its variants in the absence or presence of ZapA overexpression. Strains were grown as in (A), nascent PG was labelled with HADA as described in Material and Methods. White arrows and triangles indicate the normal and aberrant Z rings that overlap and do not overlap with a HADA signal, respectively. Note that sPG synthesis is blocked in cells expressing wild-type FtsZ but not in cells expressing FtsZ mutants. (C) Quantitation of the co-localization of ZipA-mCherry and HADA signal in (B). Data shown are the average of three experiments with more than 200 cells. Error bars indicate the standard deviation of the three experiments. Asterisks denote a significant difference based on a P value of <0.0001 (****), ns: not significant. Scale bars, 5 μm.

### ZapA cannot crosslink FtsZ mutant filaments *in vitro*

To confirm that the FtsZ mutants are resistant to the action of ZapA, we purified the FtsZ mutants and ZapA using the SUMO-tag protein purification system and examined the crosslinking activity of ZapA on FtsZ filaments by a sedimentation assay and negative stain electron microscopy. We first confirmed that the FtsZ mutants still polymerized by measuring their GTPase activity as previously described (50, 54). The K141N and V128I mutations reduced the GTPase activity by about 50%, whereas the other two single mutations did not have an obvious effect (SI Appendix, Fig. S6A). For the double and triple mutants containing the K141N mutations, their GTPase activity was about 50%-70% lower than that of wild-type FtsZ. Nonetheless, the sedimentation assay showed that all the FtsZ mutants could be pelleted in the presence of GTP and Ca^2+^ (SI Appendix, Fig. S6B), indicating that they could still polymerize. Consistent with this, negative stain electron microcopy revealed that the FtsZ mutants formed single-stranded filaments and small bundles similar to wild-type FtsZ (SI Appendix, Fig. S7). The addition of ZapA led to a significant increase in the amount of wild-type FtsZ and ZapA proteins in the pellet in the sedimentation assay (done in the absence of Ca^2+^) (Fig. 4A). Overall, ZapA had less of an effect on the amount of the FtsZ mutants in the pellet. For some mutants, especially the triple mutant, ZapA had no effect suggesting that ZapA could not crosslink these mutant FtsZ filaments effectively. In agreement with this, negative stain electron microcopy showed that ZapA promoted the formation of large bundles of wild-type FtsZ filaments, but was unable to do so with the FtsZ mutants (Fig. 4B, SI Appendix, Fig. S7). Thus, the FtsZ mutations provide resistance to the crosslinking activity of ZapA to various degrees, likely by reducing the binding affinity for ZapA.

**Fig. 4.**
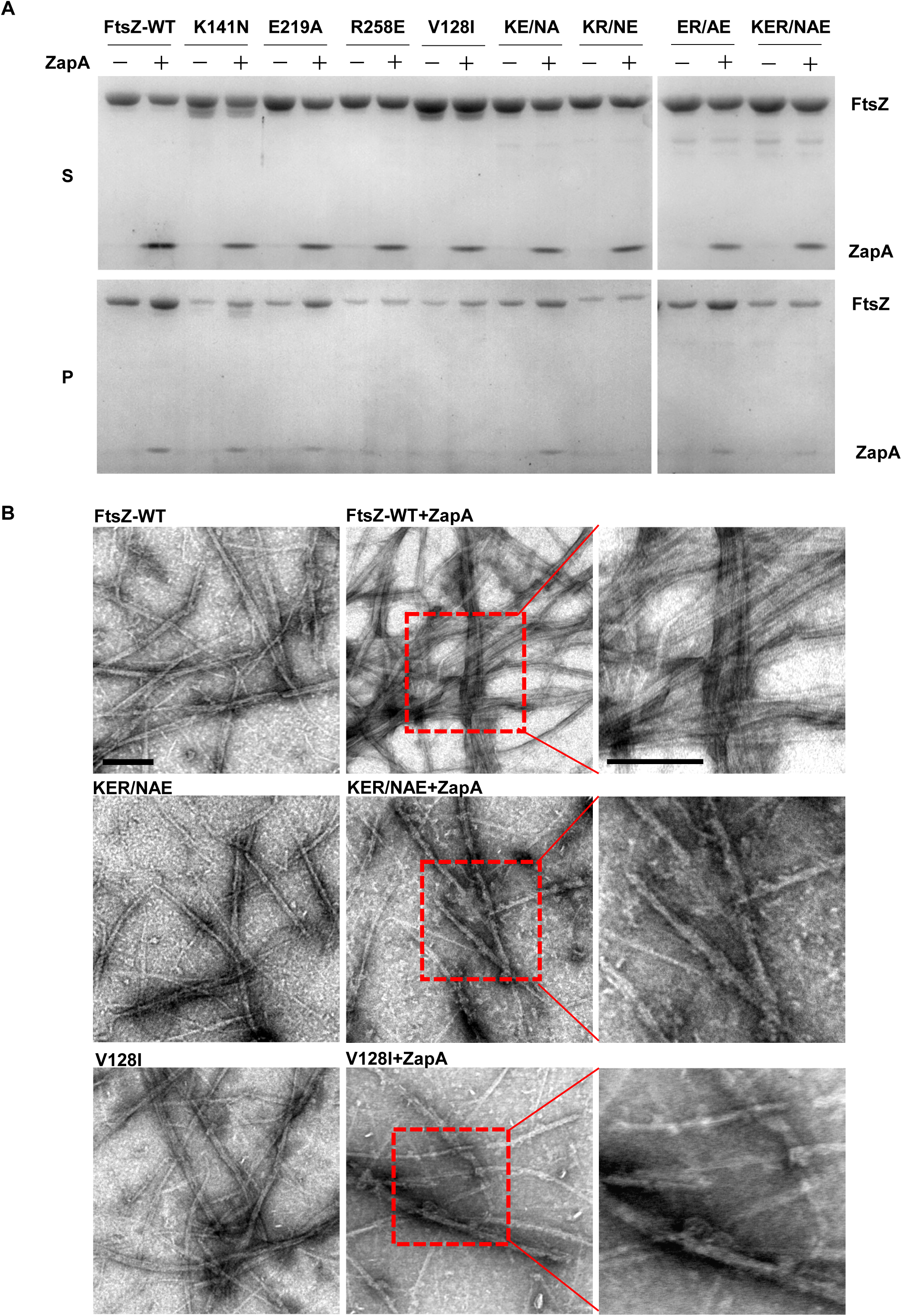
FtsZ mutants display resistance to ZapA *in vitro*. (A) Sedimentation assay to test the influence of FtsZ mutations on the crosslinking of FtsZ filaments by ZapA. FtsZ or its variants (5 μM) were mixed with or without equal molar ZapA in polymerization buffer (50 mM HEPES pH 6.8, 10 mM MgCl_2_, 200 mM KCl) in the presence of GTP (2.5 mM) in a 50 μL reaction volume. The samples were incubated at room temperature for 5 min before being centrifuged. The pellets and supernatants were analyzed by SDS-PAGE. (B) Negative stain electron microscopy analysis of the effect of the FtsZ mutations on the crosslinking of FtsZ filaments by ZapA. The reactions were performed as in (A), but the final concentration of proteins were lowered to 2.5 µM. GTP was added to a final concentration of 1 mM. Scale bars, 0.2 μm.

### FtsZ mutations weaken the interaction between FtsZ and ZapA

Next, we examined the effect of the mutations on the interaction between FtsZ and ZapA *in vivo* using the Bacterial Two-Hybrid assay (55). Since all the mutated residues are located in the globular domain of FtsZ, we constructed a truncated form of FtsZ (FtsZ^1-316^), which lacks the linker region (FtsZ^317-370^) and the C-terminal tail (FtsZ^317-383^) of FtsZ, and tested its interaction with ZapA. As shown in Fig. 5A, FtsZ^1-316^ interacted with ZapA as well as full length FtsZ, while FtsZ^317-370^ and FtsZ^317-383^ showed no interaction signal with ZapA, consistent with ZapA binding to the polymerization domain of FtsZ but not its linker or C-terminal tail. Moreover, when the single mutations were introduced into FtsZ^1-316^, most of them greatly reduced, or completely eliminated, the interaction with ZapA (Fig. 5B). All the double and triple mutations eliminated the interaction between FtsZ^1-316^ and ZapA, indicating that the mutated residues are critical for FtsZ interaction with ZapA *in vivo*.

**Fig. 5.**
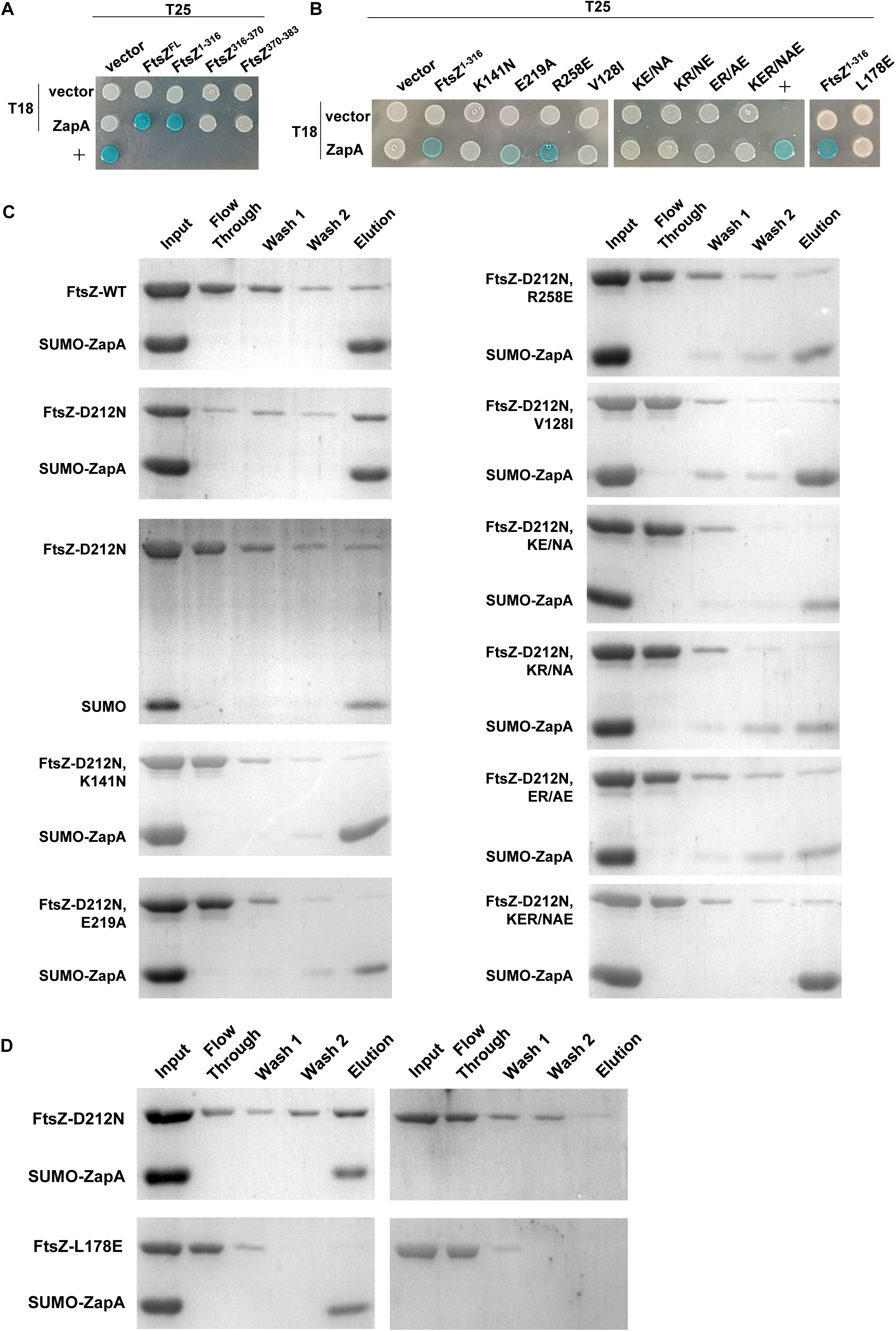
FtsZ mutations weaken the interaction between FtsZ and ZapA. (A) Bacterial two hybrid (BTH) assay to determine the domains important for FtsZ interaction with ZapA. Pairs of plasmids expressing the indicated fusions to the T18 and T25 domains of adenine cyclase were transformed into strain BTH101. A single transformant was resuspended in 1 mL of LB solution and spotted on LB plates containing IPTG-Xgal. Plates were incubated at 30°C for 12 h prior imaging. Blue indicates a positive interaction. +: positive control. A representative image from three independent experiments was shown. (B) BTH assay to access the impact of FtsZ mutations on the interaction between FtsZ and ZapA. Experiments were performed as in (A). (C) Pull-down assay to access the interaction between FtsZ mutants and ZapA. SUMO-ZapA and FtsZ or its variants were incubated and treated according to the pull-down assay described in Materials and Methods. All fractions were collected during the procedure and analyzed by SDS-PAGE. Note that since the GTPase defective mutant FtsZ^D212^ binds more readily to ZapA *in vitro*, the mutations were tested in the FtsZ^D212N^ background. (D) Pull-down assay to test the interaction between ZapA and stable FtsZ filaments (FtsZ^D212N^) or monomeric FtsZ^L178E^. Assays were performed as in (C).

To test the effect of the FtsZ mutations on the interaction between FtsZ and ZapA directly *in vitro*, we developed a pull-down assay. The SUMO tag was removed from purified SUMO-FtsZ but not from SUMO-ZapA. Since our genetic results indicated that the putative binding site for ZapA only forms when FtsZ polymerizes into filaments or oligomerizes, GTP was added into the buffers throughout the experiments. When wild-type FtsZ was incubated with SUMO-ZapA, only a small amount of the protein co-eluted with SUMO-ZapA (Fig. 5C), suggesting that wild-type FtsZ interacts weakly with SUMO-ZapA or FtsZ filaments turnover too fast to stably associate with it. If the latter was the case, then using a GTPase mutant (FtsZ^D212N^) that forms stable filaments should allow FtsZ filaments to associate more stably with ZapA. Indeed, when purified FtsZ^D212N^ was used for the pull-down assay, it was enriched in the eluate with SUMO-ZapA (Fig. 5C). This was not due to nonspecific binding of stable FtsZ^D212N^ polymers to the Ni-NTA column or the SUMO tag as most of the protein was in the flow through when it was mixed with just the SUMO tag. The establishment of the pull-down assay enabled us to test the effect of the mutations on the binding between FtsZ and ZapA after adding D212N to each of the mutants. As shown in Fig. 5C, the FtsZ mutants were not enriched in the eluate with SUMO-ZapA, indicating that the mutations weaken FtsZ’s interaction with ZapA *in vitro*. Altogether, these results indicate that the mutated residues constitute the binding site for ZapA or are important for FtsZ filaments to adopt a correct conformation to interact with it.

### ZapA prefers polymerized FtsZ over monomeric FtsZ

Based on the above results, we hypothesized that an FtsZ mutant that could not polymerize would not interact with ZapA. Previous studies demonstrated that introduction of mutations in the polymerization interface can lock FtsZ in the monomeric form, such as the L178E and L272E mutations (55, 56). As expected, introduction of the L178E mutation into the T25-FtsZ^1-316^ fusions completely eliminated the interaction between FtsZ and ZapA in the bacterial two hybrid assay (Fig. 5A). Moreover, almost all the FtsZ^L178E^ protein was found in the unbound fraction when it was mixed with SUMO-ZapA in the pull-down assay (Fig. 5D). These results strongly indicate that ZapA binds to FtsZ polymers more readily than FtsZ monomers.

### *in vivo* crosslinking indicates ZapA dimer head is in contact with the junction between FtsZ subunits in the filament

The above results strongly indicate ZapA binds to the exposed surface residues at the junction between adjacent FtsZ subunits in a filament. To confirm this, we tested if these regions of ZapA and FtsZ could be specifically crosslinked *in vivo* and if so, what effect the resistant mutations had on the crosslinking. To do this, we introduced cysteine residues at the putative interaction interface (the junction between FtsZ subunits in a filament, and the dimer head of ZapA) and tested their effects on function to see if they would be appropriate for Cys crosslinking. Through screening a number of cysteine mutation pairs in FtsZ and ZapA, we found that the pair FtsZ^N73C^ and ZapA^T50C^ was suitable for the crosslinking experiments. The N73 residue of FtsZ is located close to the putative interaction interface with ZapA (Fig. 6A). Moreover, complementation tests showed that the N73C mutation did not affect FtsZ’s ability to support cell growth, even when it was combined with the mutations affecting FtsZ’s interaction with ZapA (SI Appendix, Fig. S8A). Also, N73C did not provide resistance to ZapA overexpression, suggesting that it does not affect interaction with ZapA (SI Appendix, Fig. S8B). The T50 residue of ZapA was chosen for Cys substitution because it is located in the ZapA dimer head and adjacent to many residues found to be important for ZapA’s interaction with FtsZ (41) (Fig. 6B). The endogenous C19 residue of ZapA was also changed to alanine to avoid potential interference. A ZapA-GFP fusion containing these two mutations was still toxic indicating they did not affect the interaction with FtsZ (SI Appendix, Fig. S8C). If the introduced cysteine residues on FtsZ and ZapA were indeed located in the interaction interface, they may be cross-linked with the thiol-specific compound bismaleimidoethane (BMOE) to generate a cross-linked FtsZ-ZapA-GFP species that is expected to have a molecular weight of 80 kDa (FtsZ: 40 kDa; ZapA-GFP: 40 kDa). Indeed, we observed such a migrating species on SDS-PAGE (CLS: crosslinked species) when these cysteine containing proteins were coexpressed in the presence of BMOE (SI Appendix, Fig. S8C), suggesting that the cysteine pair was close to the intermolecular interaction interface. We also detected a band with a molecular weight about 100 kDa, which likely represents FtsZ crosslinked to an unknown interaction partner containing a cysteine or possibly through an amine group (BMOE can also react with molecules with an amine group inefficiently). Consistent with this, the non-specific cross-linked FtsZ product was also detected when FtsZ^N73C^ was expressed alone (SI Appendix, Fig. S8C). Strikingly, introduction of the mutations resistant to ZapA overexpression into FtsZ^N73C^ greatly reduced the 80 kDa crosslinked species (Fig. 6C), suggesting that they weaken the interaction between FtsZ and ZapA. For an unknown reason, the band around 100 kDa was differentially affected by the mutations, suggesting that some of the mutations might also affect FtsZ’s interaction with another unknown protein. Nonetheless, these results confirm that the dimer head of ZapA is in contact with the junction between FtsZ subunits in the filament and mutations resistant to ZapA overexpression disrupt this contact.

**Fig. 6.**
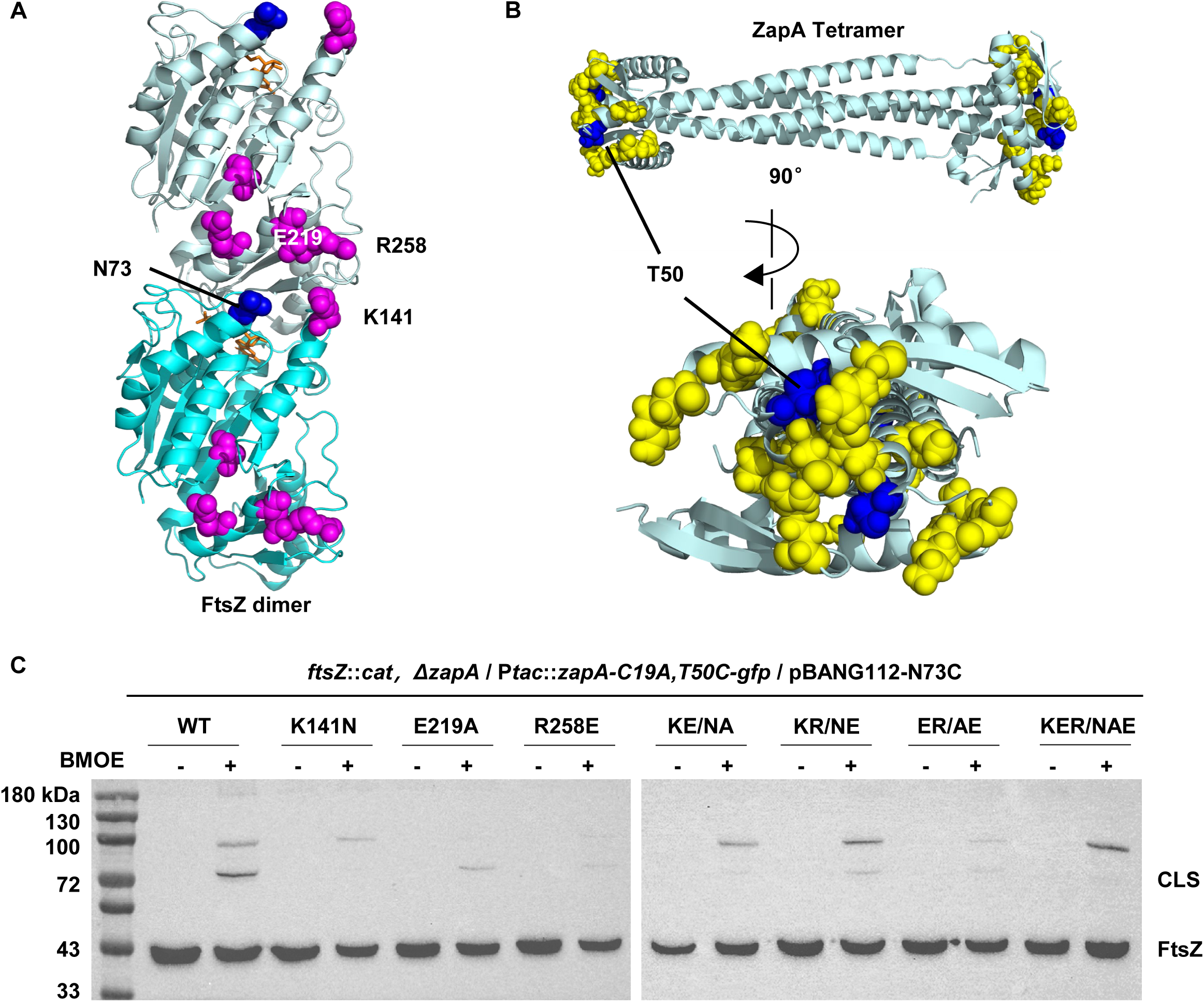
*In vivo* BMOE crosslinking assay to test the influence of the mutations on the interaction between FtsZ and ZapA. (A) Location of FtsZ mutations in an FtsZ dimer from the filament structure of *Klebsiella pneumoniae* FtsZ (PDB# 8IBN). Residues affecting FtsZ interaction with ZapA were colored magenta, the cysteine mutation N73C was colored blue. GMPCPP is shown as stick in brown. Residues number are according to *E. coli* FtsZ. (B) Location of mutations in *E. coli* ZapA tetramer structure (PDB# 4P1M). The cysteine mutation T50C was colored blue, whereas residues important for ZapA interaction with FtsZ are shown as spheres in yellow. (C) BMOE-crosslinking assay to test the effect of FtsZ mutations on FtsZ’s interaction with ZapA *in vivo*. Cells expressing FtsZ^N73C^ or its variants carrying the FtsZ mutations and ZapA^C19A,T50C^ were treated with BMOE or DMF for 15 min. Cells were then harvested by centrifugation and lysed in 1×SDS-PAGE buffer, boiled for 10 min and loaded onto SDS-PAGE gel for western blot as described in Materials and Methods.

### ZapA also binds to the N-terminal tail of FtsZ

Deep learning-based protein structure prediction approaches have been widely employed to generate protein complex structure models over the last few years. Therefore, we employed AlphaFold 3 to generate structural models of the FtsZ-ZapA complex (57). Disappointedly, the prediction did not produce any high-confidence structural models for the complex, regardless of the ratio between FtsZ and ZapA (2:2, 2:4 or 4:4). In fact, none of the structural models was able to predict the binding of ZapA to the junction between FtsZ subunits as deduced from the analysis above. Nonetheless, in all the structural models, the extreme N-terminus of FtsZ (amino acids 1-10) inserts into a groove in the dimer head of ZapA, making extensive contacts via both polar and hydrophobic interactions (Fig. 7A, SI Appendix, Fig. S9), suggesting that this motif may be also involved in the binding to ZapA. Sequence analysis of FtsZ across the bacterial kingdom revealed that this motif is quite conserved in both the Gram^+^ and Gram^-^ bacteria encoding ZapA, featured by a large hydrophobic residue (F or L) and a glutamate residue at the second and third position, respectively (Fig. 7B, SI Appendix, Fig. S10A). To test if this motif was involved in interaction with ZapA, we mutated the first 10 amino acids of FtsZ by alanine-scanning and tested whether the mutants provided resistance to ZapA overexpression toxicity. All mutants, except for the E3A substitution, was able to complement an FtsZ depletion strain (Fig. S10B), suggesting that the E3 residue is critical for FtsZ function, whereas the other residues are not. Interestingly, the FtsZ^F2A^ and FtsZ^P4A^ mutants conferred strong and weak resistance to ZapA overexpression toxicity, respectively (Fig. 7C). We therefore focused on the FtsZ^F2A^ mutant and found that Z rings (ZipA-mCherry as a proxy) formed by this mutant were not disrupted by ZapA overexpression and sPG synthesis (labelled with HADA) was not significantly hindered (Fig. 7D-F). Additionally, introduction of the F2A mutation into FtsZ^1-316^ completely eliminated its interaction with ZapA in the BTH assay (Fig. 7G). Thus, the N-terminal motif of FtsZ is indeed important for its interaction with ZapA *in vivo*.

**Fig.7.**
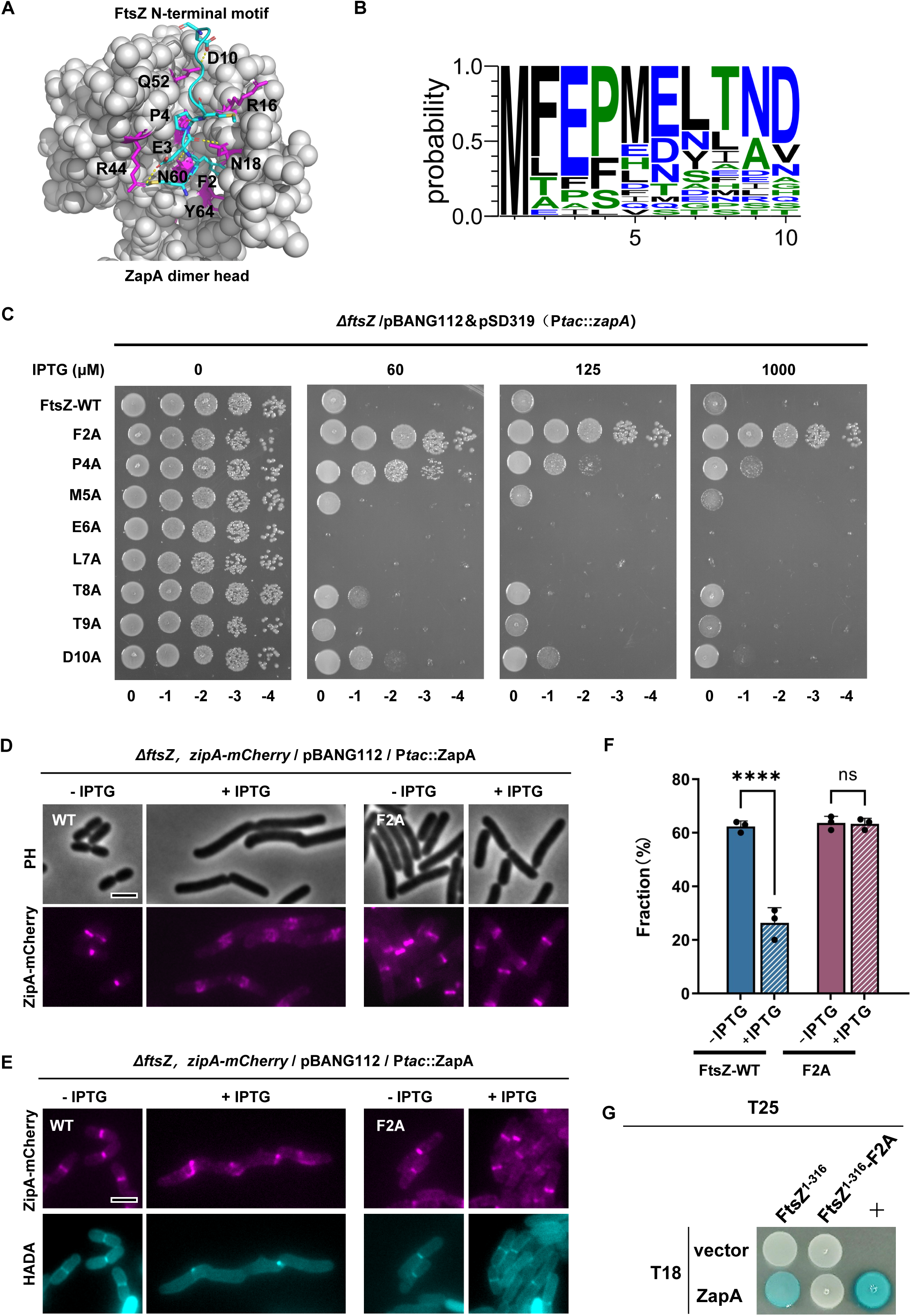
The N-terminal motif of FtsZ is important for its interaction with ZapA *in vivo*. (A) AlphaFold 3 model of the FtsZ-ZapA complex indicate that the N-terminal segment of FtsZ interacts with ZapA. The dimer head of ZapA is colored grey, while the N-terminal motif of FtsZ is colored cyan. Residues important for interaction are indicated (FtsZ: F2, E3, P4 and D10; ZapA: R16, N18, R44, Q52 and N60). (B) Sequence logo of the N-terminal motif of FtsZ across diverse bacterial species using Weblogo3. Alignment of the FtsZ N-terminal sequences was provided in SI Appendix Fig. S10A. (C) Spot test of the effect of mutations in the N-terminal motif of FtsZ on the resistance to ZapA overexpression. A plasmid expressing ZapA (pSD319) was transformed into strains expressing different FtsZ mutants, and the transformants were subjected to a spot test on plates with or without IPTG. (D) Representative images of Z rings (ZipA-mCherry) in cells expressing wild type FtsZ or FtsZ^F2A^ in the absence or presence of ZapA overexpression. (E) Representative images of co-localization of ZipA-mCherry with HADA in cells expressing wild type FtsZ or FtsZ^F2A^ in the absence or presence of ZapA overexpression. (F) Quantitation of the co-localization of ZipA-mCherry and HADA signal in (E). Data shown are the average of three experiments with more than 200 cells for each. Error bars indicate the standard deviation of three experiments. Asterisks denote a significant difference based on a P value of <0.0001 (****). ns: not significant, Scale bars, 5 μm. (G) BTH assay to test the impact of the F2A mutation on the interaction between FtsZ^1-316^ and ZapA. The test was carried out as Fig. 5A.

To confirm that the N-terminal motif of FtsZ was directly involved in the interaction between FtsZ and ZapA, we purified the FtsZ^F2A^ mutant and examined its interaction with ZapA. We confirmed that FtsZ^F2A^ polymerized by sedimentation assay (Fig. 8A) and then introduced the D212N mutation to carry out the pull-down assay. As shown in Fig. 8B, while FtsZ^D212N^ co-eluted with SUMO-ZapA, FtsZ^D212N, F2A^ did not, suggesting the F2A mutation reduces FtsZ interaction with ZapA. We also examined the effect of the F2A mutation on FtsZ’s interaction with ZapA by the sedimentation assay and negative stain electron microscopy. As shown in Fig. 8C, FtsZ^F2A^ did not sediment in the presence of ZapA. Also, while ZapA crosslinked wild-type FtsZ filaments into large bundles, less bundles of FtsZ^F2A^ filaments were observed on the grids (Fig. 8D). Moreover, the bundles formed by FtsZ^F2A^ and ZapA were much smaller and less organized in comparison to those formed by wild-type FtsZ and ZapA. Together, these results demonstrate that F2 in the N-terminal motif substantially contributes to the interaction between FtsZ and ZapA *in vitro*.

**Fig. 8.**
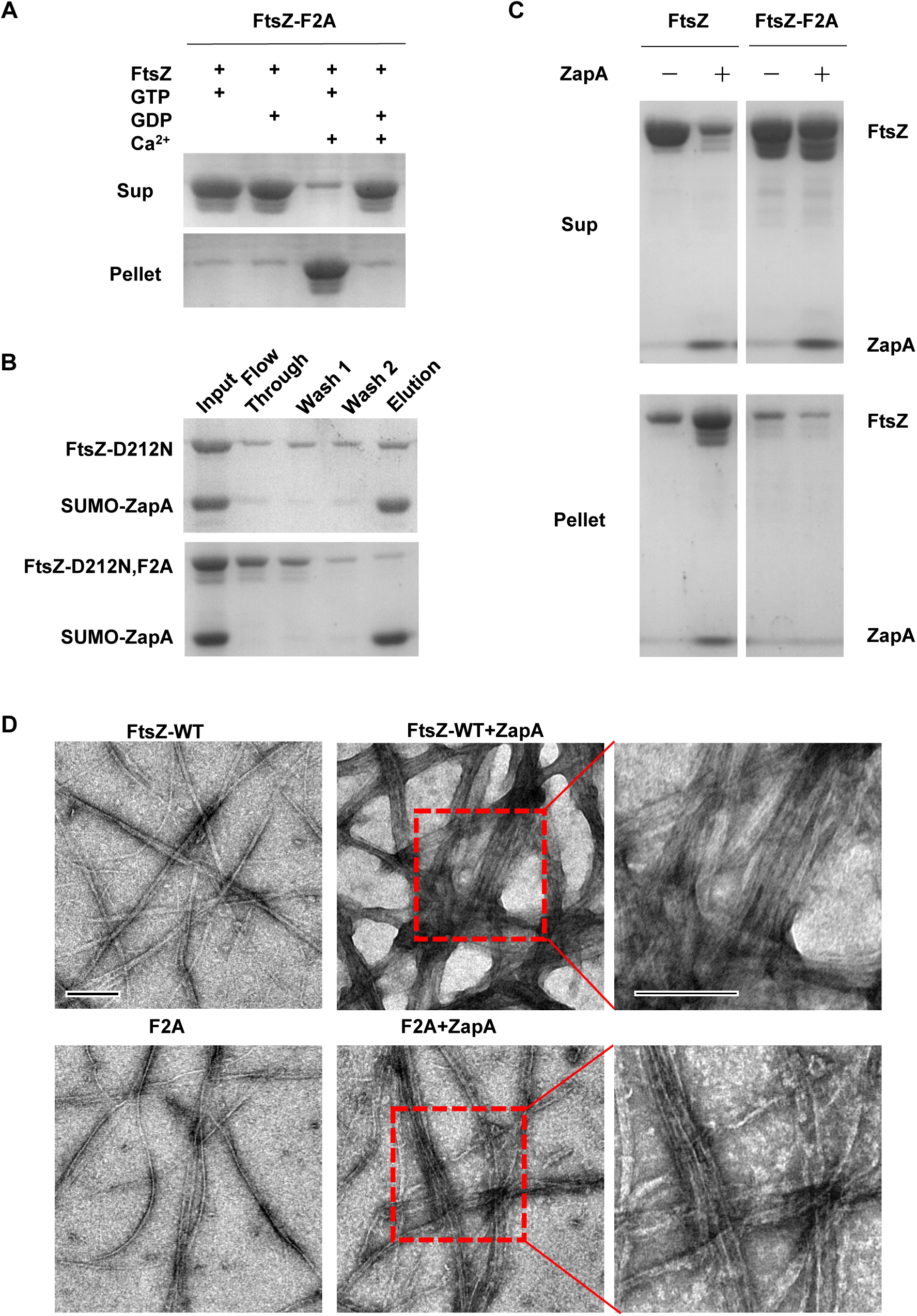
The N-terminal motif of FtsZ is important for its interaction with ZapA *in vitro*. (A) Sedimentation assay to test the polymerization of FtsZ^F2A^. FtsZ or FtsZ^F2A^ (5 μM) were mixed in polymerization buffer (50 mM HEPES pH 6.8, 10 mM MgCl_2_, 200 mM KCl) in the presence of GTP (2.5 mM) and Ca^2+^ in a 50 μL reaction volume. The samples were incubated at room temperature for 5 min before being centrifuged and the pellets and supernatants were analyzed by SDS-PAGE. (B) Pull-down assay to assess the effect of the F2A mutation on FtsZ’s interaction with ZapA. SUMO-ZapA and FtsZ^D212N^ or FtsZ^D212N, F2A^ were incubated and treated according to the pull-down assay described in Materials and Methods. All fractions were collected during the procedure and analyzed by SDS-PAGE. (C) Sedimentation assay to test the impact of the F2A mutation on FtsZ’s interaction with ZapA. The reactions were prepared as in (A) with or without equal molar of ZapA. The samples were incubated at room temperature for 5 min before being centrifuged and the pellets and supernatants were analyzed by SDS-PAGE. (D) Negative stain electron microscopy analysis of the effect of the F2A mutation on FtsZ interaction with ZapA. The reactions were performed as in (A), but the final concentration of proteins was lowered to 1 µM. GTP was added to a final concentration of 1 mM. Scale bars, 0.2 μm.

### Disruption of the interaction between ZapA and FtsZ prevents its midcell localization

Lastly, we tested if the presence of the FtsZ mutations prevented the localization of ZapA *in vivo*, since its localization depends upon its interaction with FtsZ. To do this, we introduced *zapA-gfp* into the chromosome in the strains expressing wild-type or FtsZ variants by the λ-Red recombineering system (53). We also deleted *zapB* in these strains because it interacts with ZapA to form an FtsZ-independent cloud structure at midcell by a linkage to MatP (36). As shown in Fig. 9, ZapA-GFP localized to the division site in close to 80% of cells expressing wild-type FtsZ. However, it localized to midcell in only about 40% of the cells expressing the FtsZ^F2A^ mutant, accompanied by a substantial increase in cytoplasmic fluorescence. To our surprise, the single or double mutations at the junction between FtsZ subunits only modestly reduced midcell localization of ZapA-GFP. Nonetheless, we did observe a strong reduction of ZapA-GFP midcell localization in the triple mutant (K141N, E219A and R258E). Although mutations at the N-terminal motif or at the junction between FtsZ subunits provide strong resistance to ZapA overexpression, none of them completely blocked the midcell localization of ZapA-GFP. We thus combined the F2A mutation with the mutations at the junction and tested if their effects on the localization of ZapA-GFP were additive. As expected, the addition of any mutation at the junction to the F2A mutant further reduced the midcell localization of ZapA-GFP and increased its cytoplasmic localization (Fig. 9). More importantly, a combination of the triple mutations at the junction and the F2A mutation almost completely eliminated the midcell localization of ZapA-GFP. Thus, both the N-terminal motif and the junction between FtsZ subunits in the filaments are important for ZapA binding to FtsZ *in vivo*, and disruption of both binding sites is necessary to eliminate their colocalization.

**Fig. 9.**
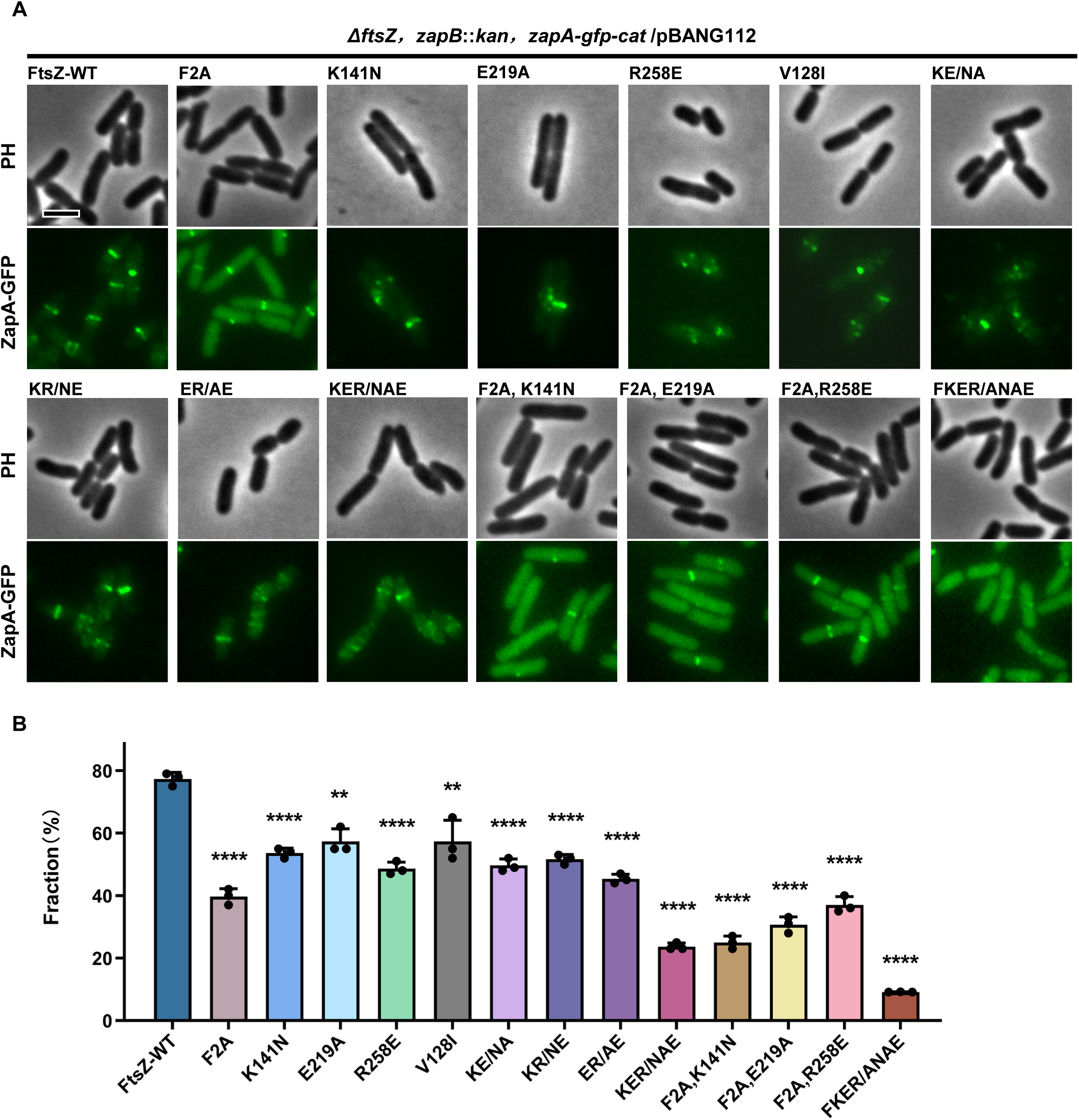
Localization of ZapA depends on its interaction with FtsZ. (A) Representative images of ZapA localization in cells expressing FtsZ mutants. Exponentially growing cultures of CYb1 (W3110, ftsZ^0^ *zapA-gfp-cat zapB::kan* /pACYC, *ftsZ*) and derivatives carrying various *ftsZ* alleles were visualized by fluorescence microscopy to assess ZapA-GFP localization at 30°C. Scale bars, 5 μm. (B) Quantification of the localization of ZapA-GFP in (A). Data are present as mean value ± s.d. Asterisks denote a significant difference based on a P value of <0.0001 (****), 0.0008< P <0.0016 (**). Number of cells analyzed (WT: n=334; F2A: n=836; K141N: n=275; E219A: n=352; R258E: n=239; V128I: n=242; KE/NA: n=289; KR/KE: n=347; KER/NAE: n=876; F2A,K141N: n=475; F2A,E219A: n=918; F2A,R258E: n=542; FKER/ANAE: n=943). Scale bar, 5 μm

## Discussion

ZapA is a very widely conserved cell division protein that facilitates Z ring assembly in diverse bacteria, but the mechanism remains unclear, in part because the binding site on FtsZ had not been determined. In this study, we found that mutations at the junction between adjacent FtsZ subunits within filaments and in an N-terminal motif of FtsZ substantially reduced its interaction with ZapA, indicating that ZapA binds to both these regions of FtsZ. This dual binding mode enables ZapA to employ the polymerization dynamics of FtsZ and efficiently crosslink FtsZ filaments in the Z ring. Moreover, we found that mutations internal to the FtsZ molecule can also significantly affect its interaction with ZapA, suggesting that ZapA recognizes a specific conformation of the filament. Taken together, our results suggest a model in which ZapA tetramers grab the N-terminal tails of FtsZ to bind to the junctions between FtsZ subunits in filaments such that they can straighten the longitudinal interaction of FtsZ like a staple and simultaneously crosslink FtsZ filaments in a parallel orientation.

The mechanism by which ZapA promotes Z ring assembly has been extensively investigated *in vivo* and *in vitro*. Early studies suggested that ZapA facilitates Z ring formation by crosslinking individual FtsZ filaments (20, 58), but subsequent studies indicated that ZapA may be more important in aligning FtsZ clusters containing multiple filaments into condensed Z rings (21). A recent *in vitro* reconstitution study of the effect of ZapA on the dynamic behavior of membrane bound FtsZ filaments provided important insight into its mechanism. It was shown that membrane-bound FtsZ filament bundles self-organize into swirling ring-like structures which reorganize into elongated bundles after the addition of ZapA (46). Using high-resolution fluorescence microscopy and quantitative image analysis, the authors showed that ZapA aligned FtsZ filaments in parallel, leading to the straightening and stabilization of the filament bundles, thereby the collapse of the swirling ring-like structures (46). However, ZapA only binds to FtsZ filaments transiently and has no effect on filament length or treadmilling velocity (46). In line with these observations, we found that overexpression of ZapA *in vivo* did not affect the treadmilling speed of FtsZ filaments but altered their treadmilling orientation. Presumably, the proper curvature and orientation of FtsZ filaments are disrupted by excess ZapA so that they could not align properly into mature Z rings. Notably, these FtsZ filaments were still able to accumulate at potential division sites as disorganized structures in the filamentous cells, suggesting that they were still responsive to the regulation of the spatial regulators. These *in vitro* and *in vivo* observations are consistent with the idea that ZapA organizes the FtsZ filament network in the Z ring without affecting the dynamics of individual FtsZ filaments.

The crystal structures of ZapA showed that it forms an elongated anti-parallel tetramer with two dimer heads connected by their C-terminal coiled-coils (40, 41). The ZapA tetramer is able to crosslink FtsZ filaments into ladder-like structures or large bundles with each dimer head bound to a filament (20, 40, 45). It may also stabilize the longitudinal interaction between FtsZ subunits within the filaments (20, 58). However, how exactly ZapA interacted with FtsZ to achieve these functions was unknown. A previous study tried to map the contact sites between ZapA and FtsZ *in vitro* by using chemical cross-linking coupled with mass spectrometry. They found that residues K51 and K66 of FtsZ were crosslinked to the K42 residue of ZapA (44). Based on this crosslinking information and mutational analysis of ZapA, a model for the complex of ZapA tetramer and FtsZ filaments was generated by docking, in which the ZapA dimer head binds to a region close to the polymerization interface of FtsZ (44). However, this model is likely rudimentary because the crosslinking information is very limited.

In this study, we found that mutations altering residues in the junction between FtsZ subunits or internal to FtsZ confer resistance to ZapA overexpression, including K66, V128, K140, K141, V193, R214, T215, S218, E219, L248, R257, R258, N289 and T291. Notably, the K66 residue was identified in the previous crosslinking experiment (44). Through in-depth characterization of three representative residues in the junction region (K141, E219, R258) by a combination of genetic, biochemical and cellular approaches, we found that these mutations weaken the interaction between FtsZ and ZapA, indicating that ZapA binds to the junction between FtsZ subunits in the filaments. This indicates that the interaction interface for ZapA is only available following FtsZ polymerization. Consistent with this, we showed that ZapA bound to stable FtsZ filaments (formed by the GTPase defective FtsZ^D212N^) better than dynamic wild-type FtsZ filaments and did not bind to a monomeric mutant (FtsZ^L178E^). It is notable that a majority of the mutated residues affecting FtsZ’s interaction with ZapA are charged residues, and many previously reported residues important for ZapA to interact with FtsZ are also charged. This indicates that the interaction between the dimer head domain of ZapA and the junction of FtsZ is largely via electrostatic interactions. It is also noticeable that the deduced binding site for ZapA covers a large surface exposed area at the junction between FtsZ subunits in a filament. This may explain why single mutations at this interface are not destructive enough to block the interaction completely, especially *in vivo*. In agreement with this, none of the reported single mutations in the dimer head of ZapA completely eliminated the interaction with FtsZ (41). Also, the presence of the additional binding site for ZapA in the N-terminus of FtsZ makes it difficult to eliminate ZapA localization to the Z ring *in vivo*.

A totally unexpected finding of this study is that ZapA also binds to an N-terminal motif of FtsZ via a groove in the dimer head. This short motif, especially the conserved F at position 2 (*E. coli* numbering), is not important for FtsZ polymerization and is not present in structures of FtsZ. Also, this motif has not been implicated in interaction with any known FtsZ binding proteins in prior studies. As a result, it was largely neglected. Although AlphaFold 3 was unable to predict the interaction interface between the dimer head of ZapA and the junction between FtsZ subunits, it implied that the N-terminal motif of FtsZ was involved in binding ZapA. Indeed, we found that mutations in this short motif significantly affected FtsZ interaction with ZapA *in vivo* and *in vitro*. Moreover, sequence alignment revealed that the F residue in this motif is highly conserved in bacteria encoding ZapA but not in those without ZapA, indicating a pivotal role of this motif in their interaction. Consistent with our finding, a previous study found that a mutation in the groove of the ZapA dimer head (N60Y) abolished the interaction with FtsZ *in vivo* (36). In the structural models, the N60 residue of ZapA interacts with the F2 residue of FtsZ. Thus, it is not surprising that mutating either one of these two residues strongly reduces or eliminates the interaction between FtsZ and ZapA. A recent Cryo-EM structure of the ZapA tetramer-FtsZ filament complex shows that the ZapA dimer head binds to this N-terminal motif of FtsZ (59), further supporting our finding. Intriguingly, in this structure ZapA tetramers do not bind to the junction between FtsZ subunits in a single filament. Also, each dimer head binds to two N-terminal motifs, each coming from an FtsZ molecule in an anti-parallel double FtsZ filaments (59). This seems to be contrary to the current working model of ZapA in which ZapA tetramers align FtsZ filaments into a parallel manner (46). It is possible that the condition used for visualization of the ZapA tetramer-FtsZ filament complex was prone to the formation of anti-parallel FtsZ filaments, allowing ZapA to bind to the N-terminal motif of FtsZ but prevented it from binding to the junction between FtsZ subunits within the filaments. Future structural studies of ZapA and FtsZ in the absence of the N-terminal motif of FtsZ is necessary to obtain a clearer picture of the interaction interface between ZapA and FtsZ filaments.

Interestingly, several mutations conferring resistance to ZapA overexpression are buried inside the FtsZ molecule and away from the deduced binding sites. Characterization of one of these mutations, V128I, showed that it reduced the interaction between FtsZ and ZapA. How could such mutations affect the interaction between FtsZ and ZapA if they are not directly involved in the interaction? It is well documented that there are substantial conformational changes in the polymerization interface and nearby regions during FtsZ polymerization and depolymerization (4, 57, 58). It is possible that the isolated mutations cause an alteration of the conformation of FtsZ within the filament, even modestly, such that the binding affinity for ZapA is reduced drastically. Nonetheless, ZapA could still bind to the N-terminal motif so that it still localized to the midcell in cells expressing the V128I mutant as shown in Fig. 9. Further studies to look at the effects of the mutations on the polymerization interface and dynamics of FtsZ are necessary to reveal the molecular basis of these mutations.

The dual binding mode between ZapA and FtsZ filaments suggests that ZapA tetramer first grabs the N-terminal tail of an FtsZ subunit to bind to a filament and then positions its dimer heads into the junction between FtsZ subunits in the filament. This binding, analogous to a constructive staple, could stabilize the junction between FtsZ subunits, leading to the straightening of the filaments. Meanwhile, as each dimer head can bind to a filament, the ZapA tetramer is able to crosslink a pair of parallel FtsZ filaments that are related a 180° rotation around the Z-axis (Fig. 10A). To validate this model, we used the HDOCK docking algorithm (60, 61) to predict the FtsZ-ZapA complex based on the filament structure of *K. pneumoniae* FtsZ and the AlphaFold 3 structural model of ZapA tetramer in complex with the N-terminal motif of FtsZ. By constraining the distance between residue 10 and 11 of FtsZ and applying additional constraints based on known mutation sites on FtsZ that affect its interaction with ZapA, we obtained a model of FtsZ-ZapA complex compatible with our finding. In this model, each dimer head of a ZapA tetramer binds to one N-terminal motif of FtsZ (in theory, each dimer head can bind two) and the junction between FtsZ subunits so that the ZapA tetramer can crosslinks two FtsZ dimers that are not entirely parallel (Fig. 10B). It is known that the coiled coil domains of ZapA can undergo conformational change, resulting in rotation of the dimer heads. As a result, it is possible that the binding between FtsZ and the dimer heads of ZapA induces conformational changes in both FtsZ filaments and ZapA, resulting in rotation of the dimer head and aligning FtsZ filaments in parallel. Moreover, FtsZ filaments have been reported to assemble into inherently curved and twisted filaments (16, 62–64), ZapA tetramers are able to crosslink multiple parallel twisted FtsZ filaments along their length into large bundles as observed under the electron microscope.

**Fig. 10.**
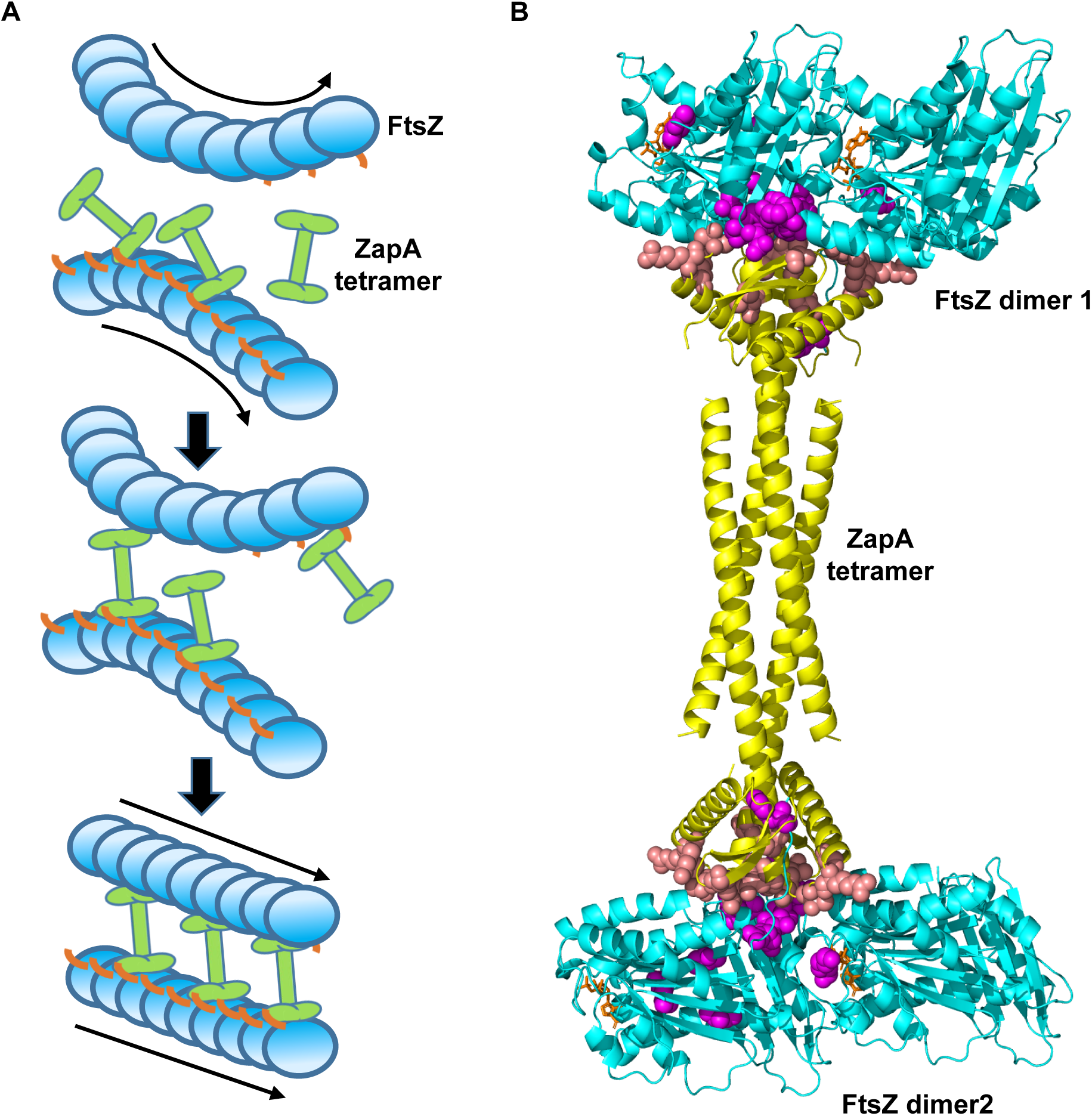
A proposed model for the mechanism of ZapA-mediated straightening and crosslinking of FtsZ filaments. (A) A diagram depicting the mechanism of ZapA. ZapA tetramers bind to the N-terminal tail of FtsZ to associate with curved FtsZ filaments. Subsequently, each dimer head of a ZapA tetramer binds to the junction of FtsZ subunits in a filament, functioning as a staple to straighten the longitudinal interaction between FtsZ subunits. Since a ZapA tetramer is bipolar with a triple 2-fold symmetry, the two dimer heads can cross-link two adjacent parallel FtsZ filaments that rotate 180° to each other. In this way, ZapA tetramers straighten and crosslink FtsZ filaments simultaneously to facilitate their organization into the Z ring. The linker and the conserved C-terminal peptide of FtsZ are omitted for simplicity. The treadmilling directions of FtsZ filaments are indicated by the arrows. (B) A docking model for the FtsZ-ZapA complex. FtsZ dimers from the filament structure of *K. pneumoniae* FtsZ (PDB# 8IBN) and ZapA tetramer in complex with the N-terminal motif of FtsZ (predicted by AlphaFold 3) were employed to generate the model by HDOCK. ZapA tetramer, colored grey, binds to both the N-terminal motif and the junction between FtsZ subunits in the filament. Residues important for FtsZ binding to ZapA are colored magenta, whereas those important for ZapA binding to FtsZ are colored pink. GTP is colored brown and shown as stick.

How could ZapA straighten and crosslink FtsZ filaments without affecting the dynamics of individual filaments? It is well documented that FtsZ filaments are semiflexible and intrinsically curved because the relative positions of adjacent FtsZ subunits in the filament are dynamically governed by its GTPase activity (65–68). The nucleotide (GTP/GDP) between FtsZ subunits governs the interaction strength and conformation of the polymerization interface (junction), with GTP stabilizing the interaction, while GTP hydrolysis weakening it. Since FtsZ filaments grow from one end and shrink from the opposite end following GTP hydrolysis, FtsZ subunits near the growing end are likely bound with GTP, whereas those near the shrinking end are likely bound with GDP. This asymmetric distribution of GTP/GDP along the FtsZ filament indicates that the junctions near the growing end are tighter while those near the shrinking end are weaker. ZapA may prefer junctions near the growing end more than those near the shrinking end. As the GTP is hydrolyzed, the configuration of the junction is altered, and ZapA dissociates from the filaments near the depolymerizing end of the filament. Although ZapA tetramer binds the junction between FtsZ subunits, accumulating evidence suggests its binding does not affect the intrinsic GTPase activity of FtsZ (46). Thus, by coupling its binding to FtsZ with the polymerization dynamics and structural flexibility of the FtsZ filaments, ZapA can organize FtsZ filaments dynamically without affecting the treadmilling speed (GTPase activity) or filament length.

## Materials and Methods

### Bacterial strains, plasmids and growth conditions

Strains were grown in LB (1% tryptone, 0.5% yeast extract, 0.5% NaCl) at indicated temperatures. When needed, antibiotics were used at the following concentrations: ampicillin = 100 mg/mL; spectinomycin = 25 mg/mL; kanamycin = 25 mg/mL; tetracycline = 12.5 mg/mL; and chloramphenicol = 15 mg/mL. The bacterial strains and plasmids used in this study were listed in SI Appendix Table S1.

### Construction of strains and plasmids

Strain CYa35 (*ftsZ^0^/pACYC, ftsZ & pEXT22, P_tac_::zapA, zipA-mCherry-spc*) was constructed in two steps. Frist, plasmid pBANG112 and plasmid pSD319 were introduced into the FtsZ depletion strain S17/pKD3C (*ftsZ^0^ /pSC101^ts^, ftsZ*). Colonies were selected at 42°C on plates containing ampicillin, kanamycin and glucose. The transformants were then streaked on LB plates with ampicillin and kanamycin at 42°C, followed by testing for sensitivity to chloramphenicol to verify plasmid pKD3C had been cured. The resultant strain was named CYm1 (*ftsZ^0^ /pACYC, ftsZ & pEXT22, P_tac_::zapA*). The *zipA-mCherry-spc* cassette was then transduced into strain CYm1 using P1 grown on CYa13 (*zipA-mCherry-spc*). The purified transductant was checked for ZipA localization and named CYa35. Strains carrying other *ftsZ* alleles and *zipA-mCherry* were constructed similarly using derivatives of pBANG112.

Strain CYb1 (*ftsZ^0^/pACYC, ftsZ*, *zapA-gfp-cat, zapB::kan*) was constructed in several steps. First strain S17/pBANG112 (*ftsZ^0^ /pACYC, ftsZ*) was constructed similarly to the CYm1. The resultant strain was then transduced with P1 from HC261 (*zapA-gfp-cat*) and transductants were selected on LB plates containing chloramphenicol. The purified transductant was named CYa1 (*ftsZ^0^, zapA-gfp-cat /pACYC, ftsZ*). Finally, the *zapB::kan* allele from SD210 (W3110, *zapB::kan*) was transduced into CYa1 by selecting for kanamycin resistance on LB plates. The resultant transductant was sequenced and named CYb1 (*ftsZ^0^,zapA-gfp-cat, zapB::kan /pACYC, ftsZ*). Strains carrying other *ftsZ* alleles and expressing *zapA-gfp* were constructed similarly using derivatives of pBANG112.

Strains for in vivo crosslinking experiments were constructed in two steps, exemplified by CYm141 (*ftsZ::cat, zapA<>frt /pACYC, ftsZ^N73C^*). First, plasmid pBC1 (*pACYC, ftsZ^N73C^*) was introduced into the strain SH19 (W3310, *zapA<>frt*). Colonies were selected on plates containing ampicillin. Second, the *ftsZ::cat* allele from CYm102 was transduced into SH19/pBC1 by selecting for chloramphenicol resistance on LB plates. The resultant transductant was sequenced and named CYm141. Strains expressing other *ftsZ* alleles were constructed similarly using derivatives of pBANG112.

Strains CYm142 (W3110 */pZH509-FtsZ-linker5-mNG*) was constructed by introducing pSY3 (*pZH509-FtsZ-linker5-mNG*) into the strain W3110. Colonies were selected on plates containing ampicillin. The purified transformant was named CYm142. Plasmid pSD319 (*pEXT22, P_tac_::zapA)* was introduced into the strain CYm142. Colonies were selected on plates containing kanamycin. The resultant transformant was named CYm143.

Plasmid pBC1 was constructed by site-directed mutagenesis using plasmid pBANG112 as the template and primer pairs FtsZ-N73C-F/R. Based on complementation tests, the cysteine substitution did not affect the function of FtsZ.

Plasmid pT18A was constructed by ligation of a BamHI/KpnI digested DNA fragment carrying the *zapA* coding sequence into pUT18C digested with the same enzymes. The DNA fragment was amplified from W3110 using primers T18-ZapA-F and T18-ZapA-R.

Plasmid pCY52 was constructed by ligation of a BsaI/XbaI digested DNA fragment carrying the *zapA* coding sequence into pE-SUMO-Amp digested with BsaI. The DNA fragment was amplified from W3110 using primers BsaI-ZapA-F and XbaI-ZapA-R.

Plasmid pCY54 and pCY55 were constructed by site-directed mutagenesis using plasmid pE-SUMO-FtsZ as the template and primer pairs FtsZ-L178E-F/R and ftsZ-D212N-F/R. These primers were listed in Table S5.

Plasmid pCY83 was constructed by ligation of a HindIII/XbaI digested DNA fragment carrying the *ftsZ^1-316^*coding sequence into pKNT25 digested with the same enzymes. The DNA fragment was amplified from pZT25 using primers pZT25-F and 316-XbaI-R.

Plasmid pCY86 was constructed by ligation of a HindIII/XbaI digested DNA fragment carrying the *ftsZ^317-383^* coding sequence into pKNT25 digested with the same enzymes. The DNA fragment was amplified from pZT25 using primers HindIII-317-F and 383-XbaI-R.

Plasmid pCY170 was constructed by replacing the *zapA* coding sequence of pSD319 with the coding sequence of *zapA-gfp*. To do this, the coding sequence of *zapA* was amplified from W3110 using primer EcoRI-ZapA-F and ZapA-GFP-R. The *gfp* fragment was amplified from pDSW210 using primer ZapA-GFP-F and GFP-HindIII-R. Overlap PCR used the purified PCR products as the template and primer pairs EcoRI-ZapA-F and GFP-HindIII-R. The resulting PCR fragment contains a fused *zapA-gfp* coding sequence, and was digested with EcoRI /HindIII and ligated into pSD319 cut with same enzymes.

Plasmid pCY173 was constructed by site-directed mutagenesis using plasmid pCY170 as the template and primer pairs ZapA-C19A-F/R and ZapA-T50C-F/R.

For plasmid pCY205 the coding sequence of *gfp-ftsI* was amplified from EC436 using primer 509-GFPU-F and FtsI-C-R. The *cat* fragment was amplified from pBAD33 using primer FtsI-C-F and C-509U-R. Overlap PCR was then used and the purified PCR products as the template with primer pairs 509-GFPU-F and C-509U-R. The resulting PCR fragment contains *gfp-ftsI* and *cat* coding sequence, and was ligated into BamHI/KpnI digested pSY3 by U-clone.

Plasmid pSY3 was constructed by seamless cloning using ClonExpress II One Step Cloning Kit (Vazyme). To generate this plasmid, the *ftsW* gene and linker (GGGGSPAPAPGGGGS) were replaced by the *ftsZ* gene with a linker (GRIGKIHIRR) from the plasmid pYD028 (pZH509-*ftsW-mNG*) (lab stock, modified from pZH509-*gfp*, a gift from Dr. Zach Hensel) (69). The *ftsZ* gene was amplified from the plasmid pXY027 (pCH-*ftsZ-gfp*) (34) using primers oSY3 and oSY4, and the vector backbone was amplified from plasmid pYD028 using primers oSY1 and oSY2. These fragments were then joined to generate plasmid pSY3.

Derivatives of pBANG112, pBC1, pCY55, pCY83, pE-SUMO-FtsZ and pZT25 carrying different *ftsZ* mutations were created by site-directed mutagenesis using the primers listed in Table S5.

### Creation of the FtsZ mutant library and selection for ZapA overexpression resistant *ftsZ* mutations

PCR random mutagenesis was used to introduce random mutations into the coding region of *ftsZ* using pBANG112 as the template and primers: 5’-CCG**GAATTC**TTCGCGGTAAATACC and 5’-GCA**TCGGC**CGGGAAATCTAC. The purified PCR fragments were then digested with EcoRI and EagI and ligated into pBANG112 digested with the same enzymes. The ligation product was then transformed into JS238 and transformants selected at 37°C on LB plates with ampicillin. All colonies that grew were pooled together and part of the pooled culture was subjected to plasmid extraction to make a stock of the FtsZ mutant library. To select for the ZapA overexpression resistant FtsZ mutants, the plasmids containing mutant *ftsZ* gene and pSD319 were co-transformed into S17/pKD3C. Colonies resistant to overexpression ZapA were selected at 42°C on plates containing ampicillin, kanamycin and 30 μM IPTG. Plasmids were isolated from the colonies that grew up and the *ftsZ* gene in the plasmids was sequenced to identify the mutations.

### Complementation test

Plasmid pBANG112 or its derivatives were used for complementation tests of the FtsZ depletion strain. Plasmids carrying the *ftsZ* alleles were transduced into the depletion strain S17/pKD3C (*ftsZ^0^ /pSC101^ts^, ftsZ*) at 30°C. Since pKD3C cannot replicate at temperature above 37°C, the complementation tests were done at 42°C by spot test.

### Fluorescence microscopy

Phase contrast and epifluorescence images were collected on an Olympus BX53 upright microscopes with a Retiga R1 camera from QImaging, a CoolLED pE-4000 light source and a U Plan XApochromat phase contrast objective lens (100X, 1.45 numerical aperture [NA], oil immersion). Green and red fluorescence was imaged using the Chroma EGFP filter set EGFP/49002, mCherry/Texas Red filter set mCherry/49008, respectively. For microscopy, a 2 μL sample of cells were immobilized on 2% agarose pads at room temperature, and a clean glass coverslip placed on top.

#### (1) Co-localization of ZipA-mCherry with GFP-FtsI

Overnight culture of CYa35 (*zipA-mCherry-spc, ftsZ^0^/pACYC::ftsZ & pEXT22, P_tac_::zapA*) carrying plasmid pCY205 (*pZH509, gfp-ftsI,cat*) was diluted 1:100 in fresh LB. After incubation at 37°C with shaking for 2.5 hours, the culture was diluted 1:10 in fresh LB then grown at 37°C. The cultures were grown for another 1.5 hour with or without 500 μM IPTG. 2 μL of the cultures was spot on 2% agarose pads for photograph.

#### (2) Fluorescent D-amino acids labeling

HADA (7-hydroxycoumarin-3-carboxylic acid-D alanine) was used to label nascent peptidoglycan. A HADA stock solution was prepared in DMSO at a concentration of 25 mM and stored at -20°C before use. A final concentration of 0.25 mM of HADA was used in all the related experiments. The cell cultures were incubated with HADA at 30°C for 1 minute. The cells were then fixed with 2.6% paraformaldehyde and 0.04% pentadiol for 15 min on an ice-bath. The cells were then collected by centrifugation at 12,000 rpm for 1 min, washed with 1×PBS, and resuspended into 1×PBS for subsequent imaging by fluorescence microscopy.

#### (3) Localization of ZipA-mCherry

An overnight culture of CYa35 (*ftsZ^0^, zipA-mCherry-spc / pACYC, ftsZ& pEXT22, P_tac_::zapA*) was diluted 1:100 in fresh LB and grown at 37°C for 2.5 hours. Then the culture was diluted 1:10 in fresh LB with or without 500 μM IPTG and cultured to OD_600_ about 0.5. 2 μL of the cultures was spotted on 2% agarose pads for photograph. Strains expressing FtsZ variants were observed similarly.

#### (4) Localization of ZapA-GFP

Overnight culture of CYb1 (*ftsZ^0^*, *zapA-gfp-cat, zapB::kan/pACYC, ftsZ*) was diluted 1:100 in fresh LB and grown at 37°C for 2.5 hours, then the culture was diluted 1:10 in fresh LB and cultured to OD_600_ about 0.5. 2 μL of the cultures was spotted on 2% agarose pads for photograph. Strains expressing FtsZ variants were observed similarly.

#### (5) Time-lapse imaging

To follow the dynamics of Z ring/structures more closely, CYm142 (W3110 /*pZH509, ftsZ-linker5-mNG*) and CYm143 (W3110/*pZH509, ftsZ-linker5-mNG & pEXT22, P_tac_::zapA*) were grown to log phase as described above and observed on a High Intelligent and Sensitive SIM (HIS-SIM) P-104WT microscope. An Apo 100×/1.5 Oil objective was used for cell imaging. FtsZ-mNG localization in real time were acquired at 10 s frame rates for 10 min from a 488 nm laser. For observation of FtsZ-mNG, the exposure time and illumination intensity were 60 ms and 0.2 (W/cm2). Samples were enclosed in a confocal glass bottom dish. Images were processed and used for production of movies in Fiji.

#### (6) FtsZ treadmilling imaging

FtsZ treadmilling speed was captured using a Nikon inverted microscope equipped with a TELEDYNE CMOS (Prime BSI Express) camera and a Nikon 100X NA 1.49 TIRF objective, the same to our previous setup (19). FtsZ-mNG expressed from P*_atc_*::*ftsZ-mNeonGreen* was excited under total internal reflection fluorescence (TIRF) mode. The images were acquired every second with an exposure time of 50 milliseconds for 150 frames.

### Bacterial Two-Hybrid analysis

To detect FtsZ-ZapA interaction, appropriate plasmid pairs encoding ZapA-T18 and FtsZ^1-316^-T25 or their variants were co-transformed into BTH101. Single colonies were resuspended in 1 ml LB medium and 2.5 μl of each aliquot was spotted on LB plates containing 100 mg/mL ampicillin, 25 mg/mL kanamycin, 40 mg/mL X-gal and 20 μM IPTG. Plates were incubated at 30°C for 12 hours before analysis. Interactions between ZapA and FtsZ^1-316^ or their variants were tested similarly.

### *in vivo* BMOE Crosslinking Assays

Strains CYm141 (*ftsZ::cat zapA<>frt /pACYC, ftsZ^N73C^*) or its derivatives were transformed with pCY173 (*pEXT22, P_tac_::zapA^C19A,T50C^-gfp, Kan^r^*). Overnight cultures of transformed strains were grown at 37°C in LB with appropriate antibiotics and then diluted 1:100 into LB containing appropriate inducer and antibiotic concentrations (pBANG112: 100 μg/mL ampicillin; pCY173: 100 μM IPTG and 25 μg/mL kanamycin). Cells were grown at 37°C to exponential phase, and 2 mL culture were harvested (10,000 rpm, 2 min), washed with 2 mL PBSG (PBS with 0.1% Glycerine) for twice, and then resuspended in 1 mL PBSG with 5 mM EDTA. Crosslinking was accomplished by treating cells with 400 μM BMOE (Thermo Scientific) at 4°C for 15 min, with DMF alone serving as the negative control. Samples were treated with 2 μL β-mercaptoethanol to quench the BMOE reaction. Cells were collected (10,000 rpm 5 min), lysed by resuspension in 50 μL 1× Loading buffer containing 5% β-mercaptoethanol, and boiled for 10 min. Whole-cell lysates were analyzed by SDS-PAGE using 10% gels and transferred to polyvinylidene fluoride membranes. Membranes were immunoblotted with rabbit antiserum against FtsZ and a goat anti-rabbit antibody. The membranes were imaged under fluorescence mode in a ChemiDoc MP system (Bio-Rad).

### Protein purification

Overnight culture of BL21/pLys harboring plasmid pE-SUMO-FtsZ (*P_T7_::his-SUMO-ftsZ*) was diluted 1:100 in fresh LB with antibiotics. After incubation at 37°C with shaking for 2 hours, 500 μM IPTG was added. The culture was grown at 37°C for another 3 hours and cells were collected by centrifugation at 8,000 rpm for 10 min at 4°C. The cells were resuspended in 20 mL lysis buffer (25 mM Tris-HCl [pH 7.5], 300 mM NaCl, 0.1 mM DTT, 20 mM imidazole and 5% glycerol). The lysates were centrifuged at 10,000 rpm for 10 min at 4°C to remove cell debris. The supernatants were removed and loaded onto pre-equilibrated Ni-NTA resin (Qiagen). The column was washed twice with high salt wash buffer (25 mM Tris-HCl [pH 7.5], 500 mM NaCl, 0.1 mM DTT, 20 mM imidazole and 5% glycerol). The bound protein was eluted with elution buffer (25 mM Tris-HCl [pH 7.5], 500 mM NaCl, 0.1 mM DTT, 250 mM imidazole and 5% glycerol) and analyzed by SDS-PAGE. The fractions with large amount of proteins were pooled together and dialyzed against the dialysis buffer (25 mM Tris-HCl [pH 7.5], 300 mM NaCl, 0.1 mM DTT and 5% glycerol) overnight, and dialyzed against the fresh dialysis buffer for 6 hours next day. After dialysis, the His-SUMO tag was cleaved with purified 6×His-tagged SUMO protease (Ulp1) for 1 hour at 30°C. The released tag and protease were removed by passing it through the pre-equilibrated Ni-NTA resin. Untagged FtsZ was collected in the flow through, concentrated and stored at -80°C. All FtsZ mutant proteins were purified similarly.

The H-SUMO-ZapA was expressed and purified from BL21 (DE3)/pLys cells containing pCY52 (*P_T7_::his-SUMO-ZapA*). An overnight culture of the strain grown in LB with ampicillin (100 mg/mL) and glucose (0.2%) was diluted 1:100 into 300 mL fresh LB medium supplemented with ampicillin (100 mg/mL) and incubated at 37°C for 2.5 hours, 1 mM IPTG was added. The culture was grown at 37°C for another 2.5 hours. The subsequent procedures were similar to purification of H-SUMO-FtsZ. The His-SUMO tag of the H-SUMO-ZapA was cleaved with purified 6×His-tagged SUMO protease (Ulp1) as H-SUMO-FtsZ.

### GTPase assay

GTPase activities of FtsZ and its mutants were determined using the NADH coupled enzymatic assay (54). The reactions were carried out at room temperature in 200 µL volume using the FtsZ polymerization buffer (50 mM HEPES-NaOH pH 6.8, 50 mM KCl, 10 mM MgCl_2_), plus 1 mM PEP, 1.2 mM NADH, 5 µL PK/LDH and 2.5mM GTP.

FtsZ or its mutants were added to a final concentration of 2.5 µM. NADH depletion is directly proportional to GTP hydrolysis, thus the reactions were continuously monitored for NADH absorbance at 340 nm for 30 minutes. The data was collected and then plotted using the Prism software, and the reaction rates were calculated using the Beer-Lambert law (A= ε×c×l).

### Pull Down assay

The pull-down assay was performed at 4°C. H-SUMO-ZapA (6 μM) and FtsZ (2 μM) were mixed in a total volume of 500 μL Pol buffer (50 mM HEPES NaOH pH 6.8, 50 mM KCl and 10 mM MgCl_2_), incubated at room temperature for 10 min and then loaded into a gravity flow column with 200 μL pre-equilibrated Ni-NTA agarose. After incubation on ice for 5 min without agitation, the mixture was allowed to pass through the column by centrifugation at 2000 rpm for 30 s at 4°C. The column was then washed with 500 μL of wash buffer (25 mM Tris-HCl [pH 7.5], 500 mM NaCl, 0.1 mM DTT, 20 mM imidazole and 5% glycerol) twice. Proteins bound to the Ni-NTA beads were eluted with 500 μL of elution buffer (25 mM Tris-HCl [pH 7.5], 500 mM NaCl, 0.1 mM DTT, 250 mM imidazole and 5% glycerol). All fractions were collected during the procedure and analyzed by SDS-PAGE. All FtsZ mutant proteins were tested similarly.

### Sedimentation assay and electron microscopy

FtsZ or its variants (5 µM) were tested for polymerization before sedimentation assay. The reactions were performed in 50 µL Pol buffer (50 mM HEPES NaOH pH 6.8, 50 mM KCl and 10 mM MgCl_2_) at room temperature. FtsZ or its variants (5 µM) was mixed with or without 10 mM CaCl_2_ at room temperature for 5 min. After the addition of 2.5 mM GTP or GDP, the reactions were kept at room temperature for 5 min and then subjected to ultracentrifugation at 100,000 rpm for 15 min at 25°C in a Beckman Optima Max XP centrifuge with a TAL130 rotor (Beckman Coulter, Inc.). Supernatants and pellets were then analyzed by SDS-PAGE. In experiments in which ZapA was added, FtsZ or its variants (5 µM) was mixed with or without untagged ZapA (5 µM) at room temperature for 5 min before the addition of 2.5 mM GTP. After 5 min at room temperature the samples were handled as described for the sedimentation assay.

To visualize the effect of ZapA on the filaments of FtsZ or its mutants by electron microscopy, FtsZ or the mutant proteins (2.5 µM) and untagged ZapA (2.5 µM) were mixed together and 1mM GTP was added to initiate polymerization. After a 10 min incubation at room temperature, 15 µL samples were loaded onto glow discharged grids. The excess solution was then blotted away after another 5 min and the grid was stained with 15 µL of 1% uranyl acetate for 1 min and then blotted away. The grids were air dried for more than 12 h and imaged with a JEM-1400plus transmission electron microscope. When comparing the effect ZapA on wild-type FtsZ and FtsZ^F2A^ filaments, the final concentration of proteins was lowered to 1 µM because FtsZ^F2A^ tends to assemble into bundles by itself at higher concentration.

### FtsZ filaments treadmilling speed and orientation analysis

The FtsZ filaments (or clusters) with obverse movement along a certain orientation were selected and segmented manually. The selected frames of the movie were used for maximum projection of intensity (MPI) using Fiji (70). The angle (θ) between the orientation of the fluorescence line in the MPI image and the long-axis of the cell was measured using the Angle Tool in Fiji. The treadmilling speed was measured as previously described (19). Briefly, drift correction was applied using the HyperStackReg plugin in Fiji if needed. Every image frame in the time-lapse fluorescence movie (in TIRF mode) was denoised using the PureDenoise plugin (Florian Luisier at the Biomedical Imaging Group (BIG), EPFL, Switzerland) in Fiji with a moving average over a 3-frame window for 10 iterations, employing global estimation. Next, each movie was enlarged to a pixel size of 21.7 nm using the bicubic interpolation method in Fiji. Subsequently, every movie was processed using the kymograph plugin (https://github.com/remiberthoz/imagej-live-kymographer) with a 3-frame average for smoothing. Kymographs were then created using the kymograph plugin from a line 13 pixels wide (∼280 nm) across the Z ring. The treadmilling speeds from both the leading and trailing edge were calculated. The box diagram plotted in OriginLab software was based on data from three independent experiments.

### Docking of FtsZ and ZapA tetramer

To validate our model for the interaction between FtsZ and ZapA, we employed the HDOCK docking algorithm using available FtsZ filament structure and AlphaFold 3 generated structural models. Since the AlphaFold3 server cannot accurately predict the complex structure of FtsZ and ZapA, and the ipTM value is quite low, the results are clearly unreliable. However, based on the predictions, the N-terminal of FtsZ is likely to interact with ZapA. Therefore, we used the AlphaFold3 server to predict the structure of ZapA in complex with the N-terminal of FtsZ (residues 1-10). The resulting model showed an ipTM value of 0.57 between the N-terminal of FtsZ and ZapA, indicating a certain level of reliability. Subsequently, we used the HDOCK docking algorithm to predict the interaction structure between the polymerization domain of FtsZ (residues 11-316) and the FtsZ N-terminal-ZapA complex. During the docking process, we constrained the distance between residue 10 and 11 of FtsZ to ensure that the N-terminal motif of FtsZ connects seamlessly with the remaining portion in the docking model. Also, our data showed that mutations at residues 66, 140, 141, 214, 215, 218, 219, 258, 289, and 291 of FtsZ, which are clustered in a specific region, weaken its interaction with ZapA. Therefore, we applied these residues as constraints to filter the generated docking models, eliminating those that did not satisfy the constraints. Subsequently, we manually inspected the remaining docking models, selecting those in which the head domain of ZapA interacts well with the constrained region of FtsZ. The final selected model was used for further analysis.

## Acknowledgments and funding sources

We thank members of the Du lab, Yang lab, Huang lab, Chen lab and Lutkenhaus lab for advice and helpful discussions to carry out this study and Yazhou Shi for construction of the plasmid pSY3. We would also like to thank Dr. Martin Loose at Institute of Science and Technology Austria for suggestions. This study was supported by National Natural Science Foundation of China (grant 32270049 and 32070032, http://www.nsfc.gov.cn/) and the Fundamental Research Funds for the Central Universities (grant 2042021kf0198) to S.D.; S.H.’s research is supported by National Natural Science Foundation of China (grant: 32161133002 and 62072199). D.Y., X.W., H.H. and X.Y.’s research is supported by National Natural Science Foundation of China (grant 32270035, http://www.nsfc.gov.cn/) and Anhui Provincial Natural Science Foundation (Award 2208085MC40).

