## Supplementary Figures and Tables for "ZapA employs a two-pronged mechanism to facilitate Z ring formation in *Escherichia coli*"

**Affiliation**
1 Hubei Key Laboratory of Cell Homeostasis, College of Life Sciences, Wuhan University, Wuhan, Hubei, China
2 Key Laboratory of Polar Environment Monitoring and Public Governance (Wuhan University), Ministry of Education, China
3 Division of Life Sciences and Medicine, University of Science and Technology of China, Hefei, Anhui, China
4 School of Physics, Huazhong University of Science and Technology, Wuhan, Hubei, China
5 State Key Laboratory of Virology, College of Life Sciences, Wuhan University, Wuhan, Hubei, China
6 Department of Microbiology, Molecular Genetics and Immunology, University of Kansas Medical Center, Kansas City, Kansas, USA

*** To whom correspondence should be addressed:**
Shishen Du
Hubei Key Laboratory of Cell Homeostasis,
College of Life Sciences, Wuhan University, Wuhan, Hubei, China


### Supplementary Figures

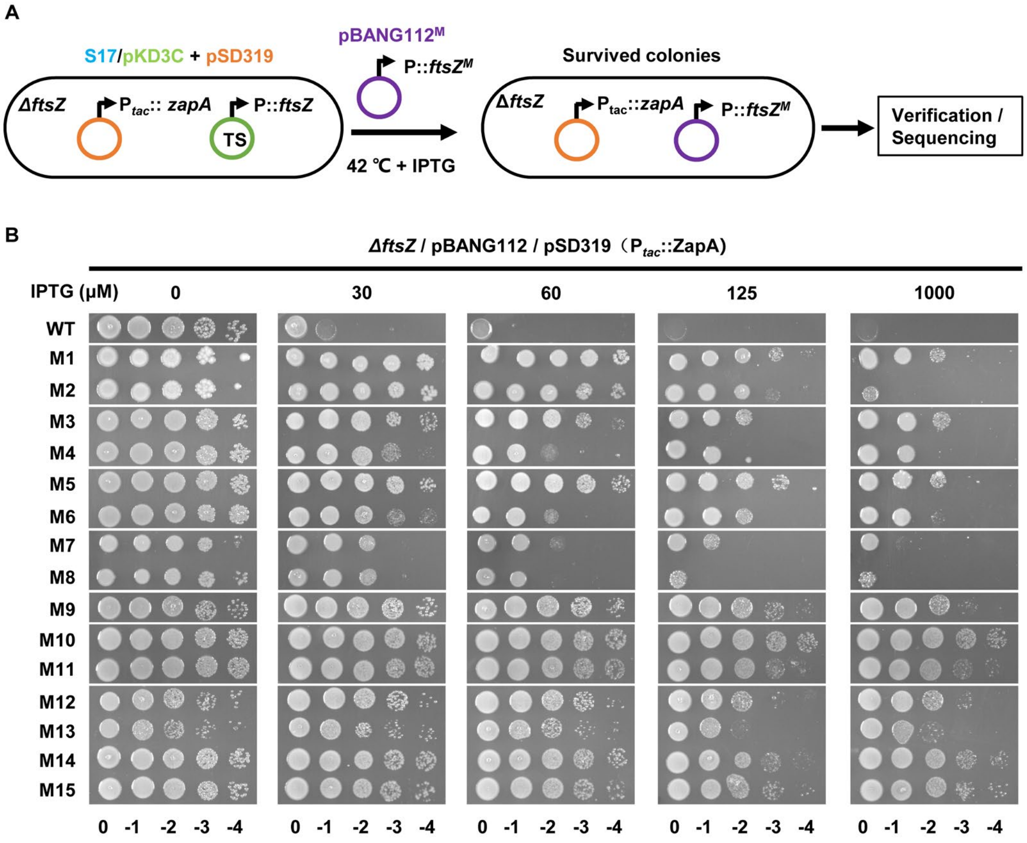


**Fig. S1 Selection for FtsZ mutants resistant to ZapA overexpression toxicity.** (A) A diagram showing the procedure for selection for FtsZ mutants. Plasmid pKD3C (in green) carrying *ftsZ* is temperature sensitive for replication, so it only complements the *ftsZ* deletion strain S17 at 30°C. The strain also harbors a plasmid expressing ZapA under an IPTG-inducible promoter (pSD319, in orange). The FtsZ mutant library ($pBANG112^M$, in purple) was introduced into the strain S17/pKD3C&pSD319 and transformants were selected in the presence of 30 μM IPTG at 42°C. Only transformants expressing functional FtsZ that provided resistance to ZapA overexpression would survive the selection. Surviving transformants were subjected to a spot test to confirm resistance to ZapA overexpression and *ftsZ* was sequenced to identify the mutations. (B) Verification of the FtsZ mutants resistant to ZapA overexpression before sequencing analysis. Transformants selected in (A) were subjected to a spot test on plates with increasing IPTG. Totally 15 mutants were isolated.

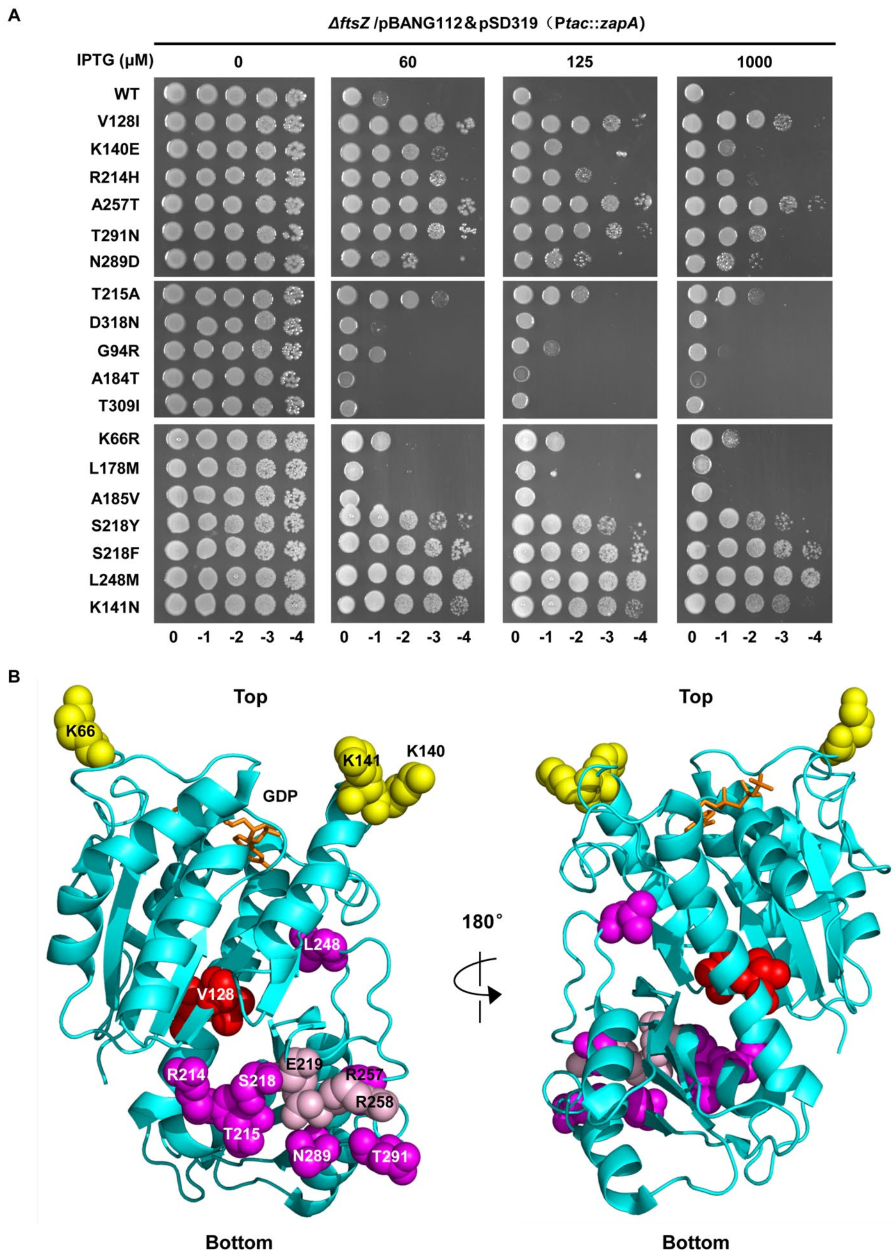


**Fig. S2 Determination of the resistance of FtsZ single substitutions to ZapA overexpression toxicity.** (A) Single substitutions identified in the 15 mutants isolated in Fig. S1B were re-introduced into pBANG112 by site-directed mutagenesis. The resultant pBANG112 derivatives carrying the single substitutions were transformed into S17/pKD3C&pSD319 and transformants selected at 42°C. Individual transformants were subjected to a spot test on plates with increasing IPTG. (B) Locations of the mutated residues in the structure of *E. coli* FtsZ (PDB# 6UNX). Residues on the surface of FtsZ are colored yellow (top face) or magenta (bottom face), whereas residues buried inside FtsZ molecule are colored red. Mutations isolated by site-directed mutagenesis are colored pink. GTP is shown as stick in brown. Residues number are indicated.

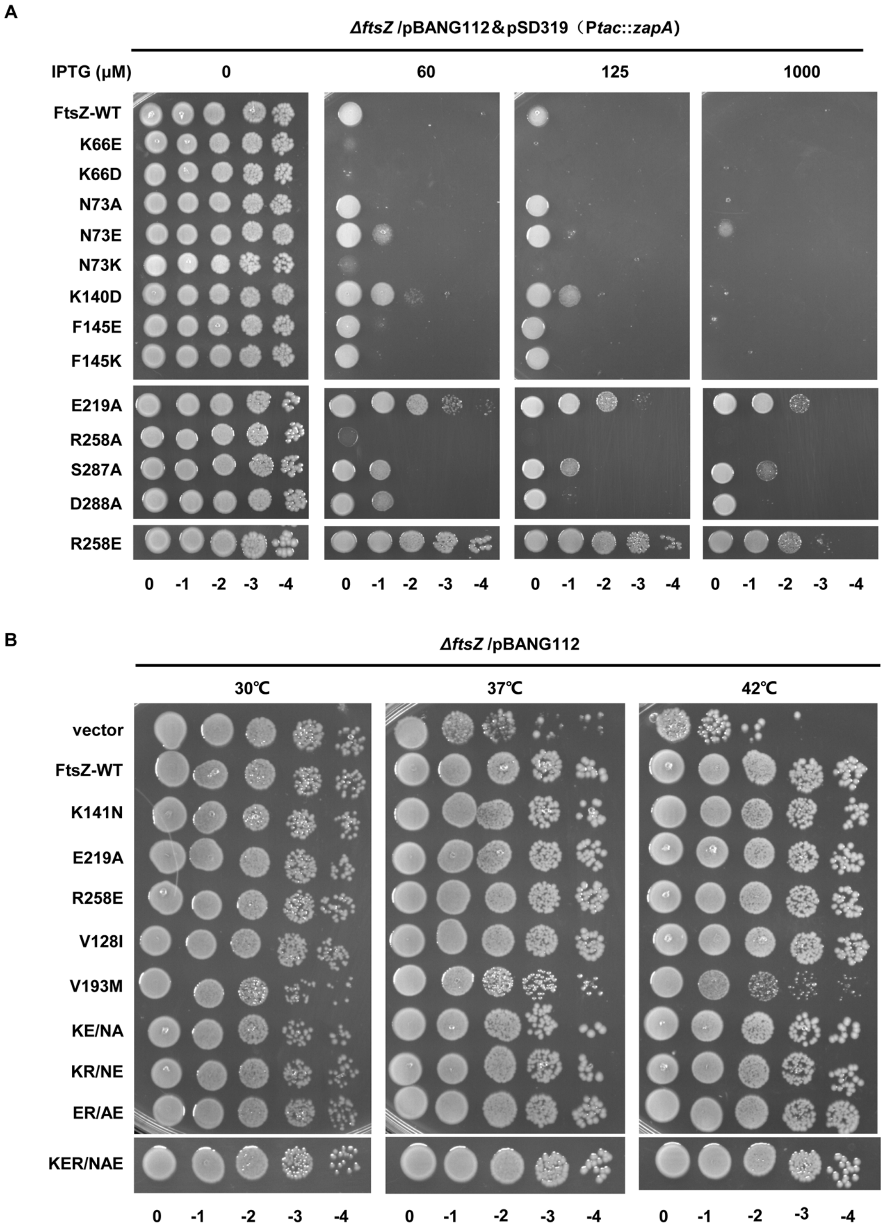


**Fig. S3 Spot test of the resistance of FtsZ mutants to ZapA overexpression and their ability to complement FtsZ depletion strain.** (A) Screening for additional FtsZ mutants providing resistance to ZapA overexpression. Residues adjacent to the mutated residues in Fig. S2 were replaced with the indicated amino acid in pBANG112. The variants were tested for resistance to ZapA overexpression as in Fig. S2A. Only E219A and R258E confer strong resistance to ZapA overexpression. (B) Complementation test of FtsZ mutants. Plasmid pKD3C carrying *ftsZ* is temperature sensitive for replication, so it only complements the ftsZ defective strain S17 at 30°C. pBANG112 or its derivatives were transformed into S17/pKD3C at 30°C. Transformants were subjected to a spot test on plates at different temperatures.

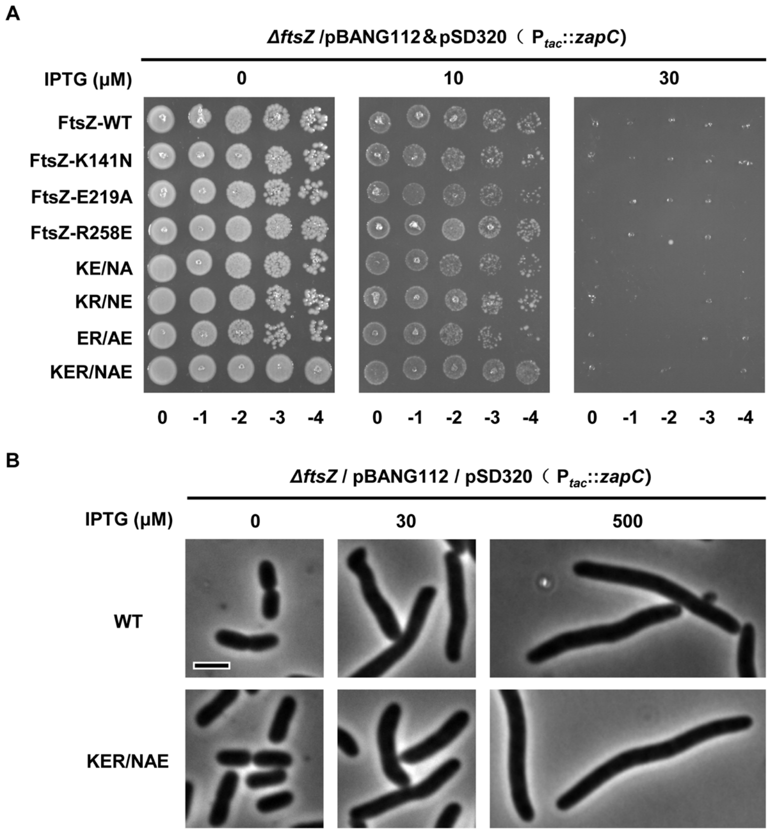


**Fig. S4 FtsZ mutants do not provide resistance to overexpression of ZapC.** (A) Spot test of the resistance of FtsZ mutants to ZapC overexpression. pBANG112 or its derivatives carrying the substitutions were transformed into S17/pKD3C&pSD320 (ZapC) and transformants selected at 42°C. Individual transformants were resuspended, serially diluted and spotted on plates with increasing concentration of IPTG at 37°C. (B) Representative image of the morphology of cells expressing FtsZ mutants and overexpressing ZapC in (A). Scale bar, 5 μm.

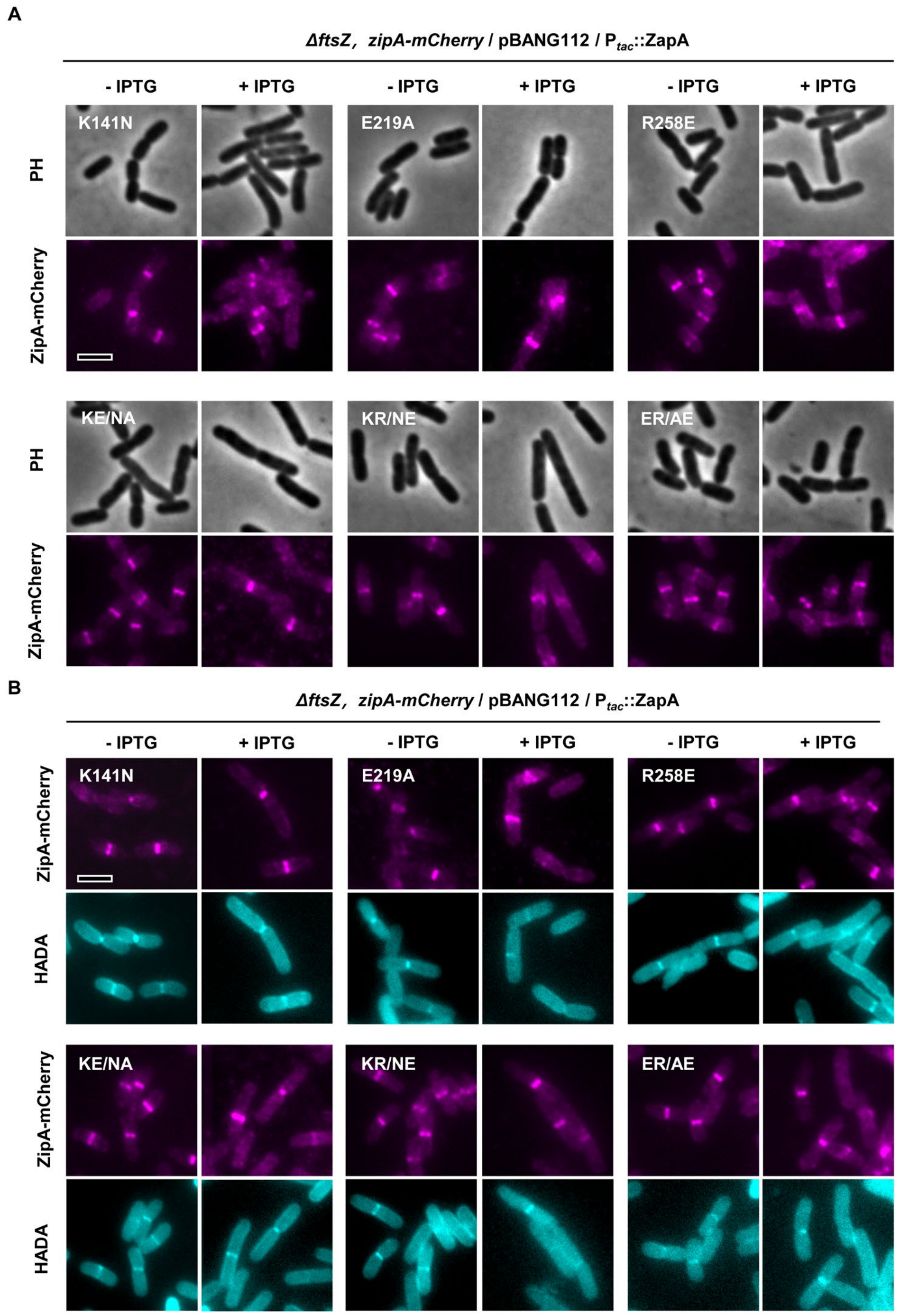


**Fig. S5 Z rings formed by FtsZ mutants are resistant to ZapA overexpression.** (A) Representative images of Z rings (ZipA-mCherry) in cells expressing FtsZ variants in the absence or presence of ZapA overexpression. Cells expressing FtsZ variants were grown in exponential phase, ZapA was induced with 500 μM IPTG, ZipA-mCherry was expressed from its native promoter. ZipA-mCherry was imaged by fluorescence microscopy. Corresponding to Fig. 3A. (B) Representative images of co-localization of ZipA-mCherry with HADA in cell expressing FtsZ variants in the absence or presence of ZapA overexpression. Strains were grown as in (A), nascent PG was labelled as described in Materials and Methods. Corresponding to Fig. 3B. Scale bars, 5 μm.

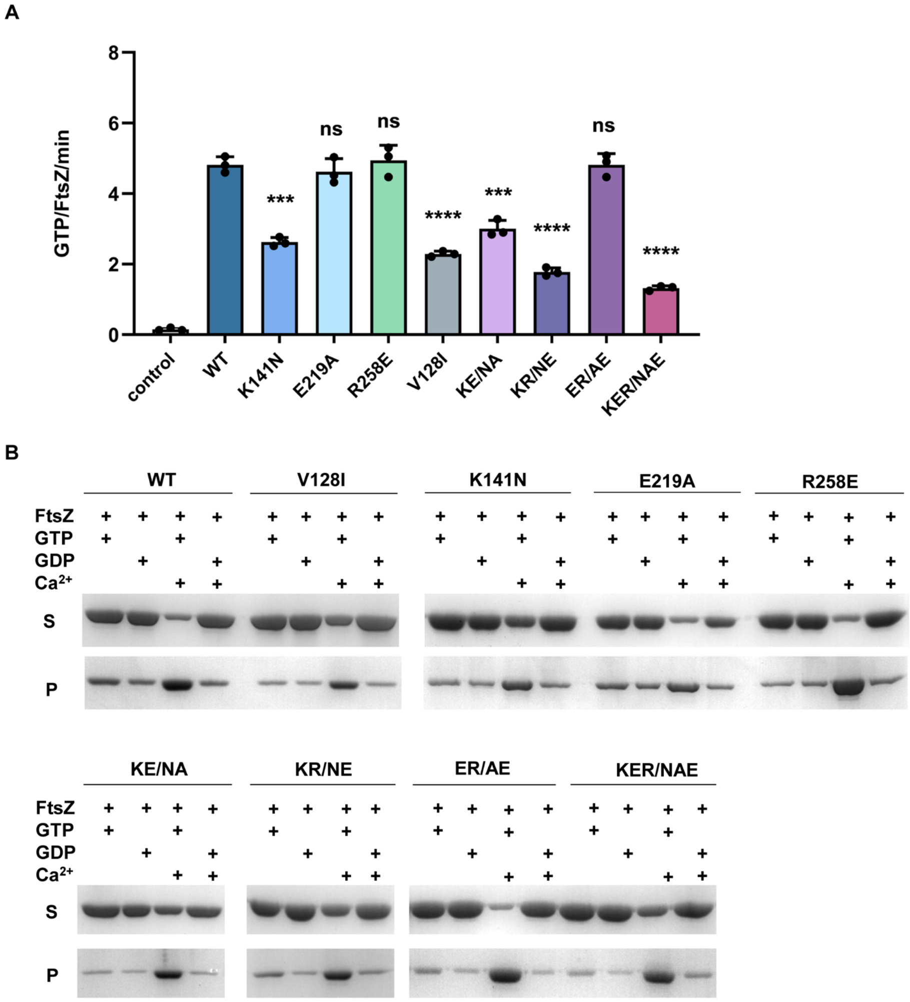


**Fig. S6 FtsZ mutants retain the ability to polymerize and hydrolyze GTP.** (A) GTPase activity of FtsZ mutants measured by the NADH coupled enzymatic assay. The reactions were carried out as described in Materials and Methods. FtsZ or the FtsZ mutants were added to a final concentration of 2.5 μM. The data was collected and then plotted using the Prism software, and the reaction rates were calculated using the Beer-Lambert law (A= ε×c×l). Asterisks denote a significant difference based on a P value of <0.0001 (****), 0.0001< P <0.008 (***), ns: not significant. (B) Sedimentation assay to access the polymerization ability of FtsZ mutants. FtsZ or its variant (5 μM) were added into polymerization buffer (50 mM HEPES pH 6.8, 10 mM $MgCl_2$, 200 mM KCl) in the presence of and GTP (2.5 mM) and 10 mM $Ca^{2+}$ in a 50 μL reaction volume. The samples were incubated at room temperature for 5 min before centrifuged and the pellets and supernatants were analyzed by SDS-PAGE. KE/NA: K141N, E219A; KR/NE: K141N, R258E; ER/AE: E219A, R258E. Scale bar, 0.2 μm.

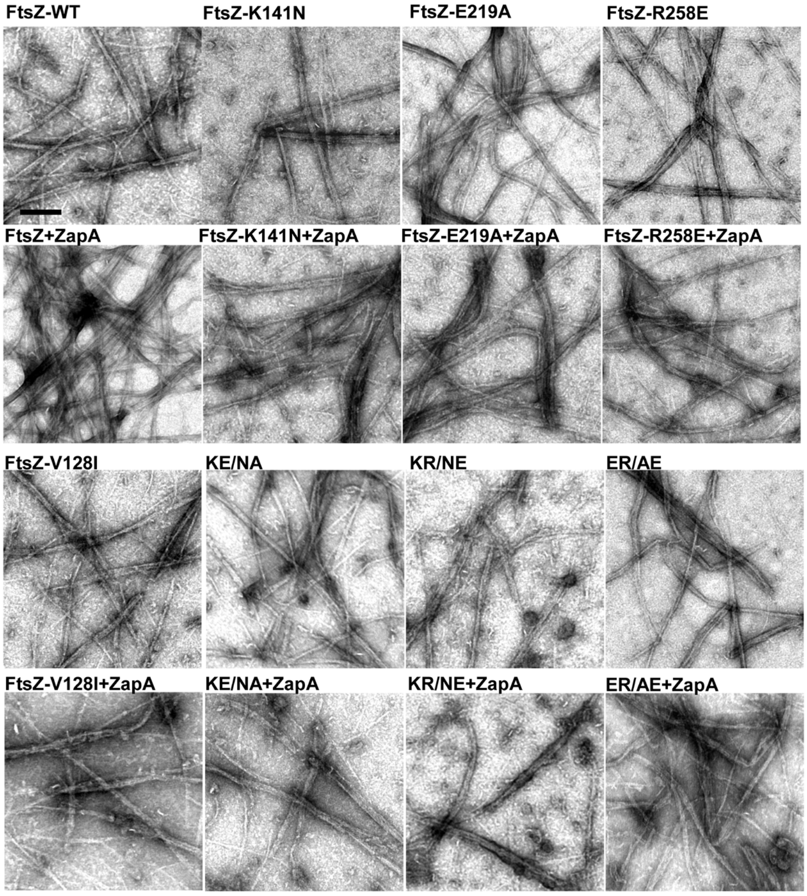


**Fig. S7 Negative stain electron microscopy analysis of the effect of FtsZ mutations on the crosslinking of FtsZ filaments by ZapA.** FtsZ and ZapA were added at a 1:1 ratio (2.5 μM) and polymerization was initiated by the addition of GTP (1 mM). After incubation at room temperature for 5 min, samples were dropped onto discharged grids and stained with 1% uranyl acetate. FtsZ filaments were examined by negative stain electron microscopy. KE/NA: K141N, E219A; KR/NE: K141N, R258E; ER/AE: E219A, R258E; Scale bar, 0.2 μm.

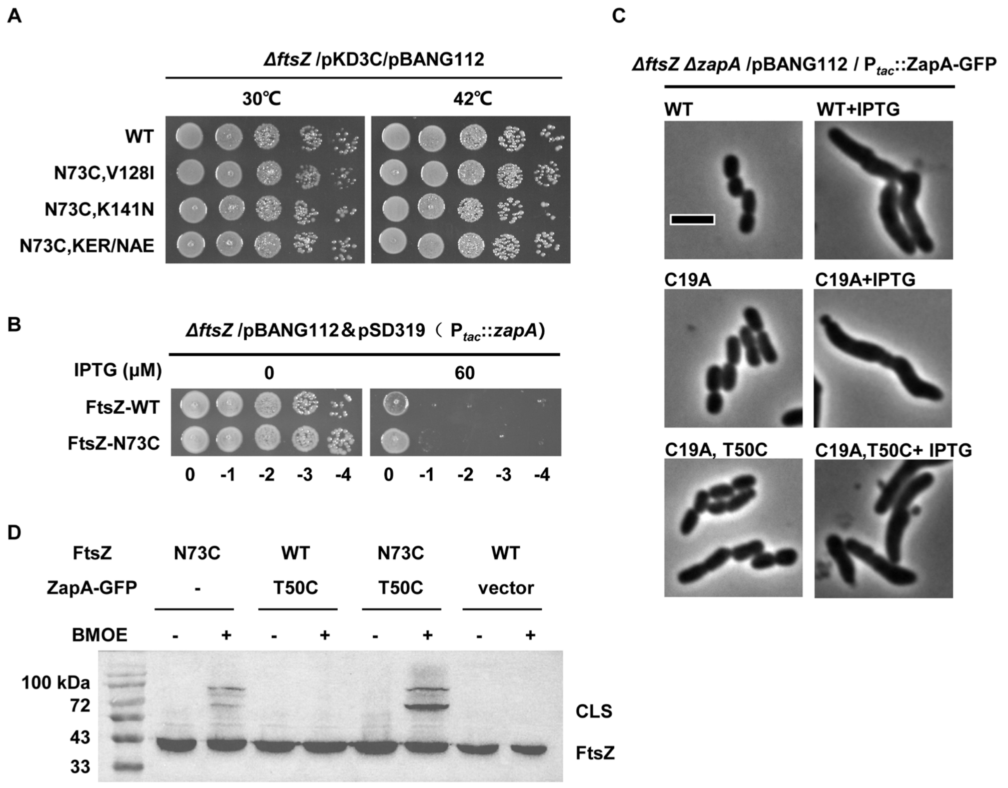


**Fig. S8 *In vivo* BMOE crosslinking assay to access the interaction between FtsZ and ZapA.** (A) Spot test to check the effect of the N73C mutation on the ability of FtsZ to complement. pBANG112 or its derivatives expressing the indicated FtsZ variants were transformed into an FtsZ depletion strain. Transformants were selected at 30°C and then subjected to a spot test at 30°C and 42°C. (B) Spot test to check the effect of N73C mutation on the sensitivity of FtsZ to ZapA overexpression. pBANG112 or its derivative expressing FtsZ$^{N73C}$ were transformed into an FtsZ depletion strain harboring a plasmid expressing ZapA. Transformants were selected at 42°C and then subjected to a spot test on plates with or without IPTG at 37°C. (C) Introduction of the T50C mutation into ZapA does not reduce its overexpression toxicity. pCY170 ($P_{tac}$::*zapA-gfp*) or its derivatives were transformed into CYm138 (*W3110, ftsZ::cam /pACYC, ftsZ*, *zapA<>frt*) . The cells were grown with or without 500 μM IPTG for 1 h and the cultures were imaged by microscopy. Scale bar, 5 μm. (D) *In vivo* BMOE crosslinking assay to test the interaction between FtsZ$^{N73C}$ and ZapA$^{(C19A)-T50C}$. Strains expressing the indicated form of FtsZ and ZapA-GFP were grown in LB in exponential phase, treated with BMOE or DMF as described in Materials and Methods. Samples were treated with β-mercaptoethanol and then cells were harvested by centrifugation and prepared for SDS-PAGE and western blot. Samples with both FtsZ and ZapA-GFP cysteine mutant pairs exhibited a significant band at 72-100 kDa corresponding to a crosslinked FtsZ-ZapA-GFP polypeptide.

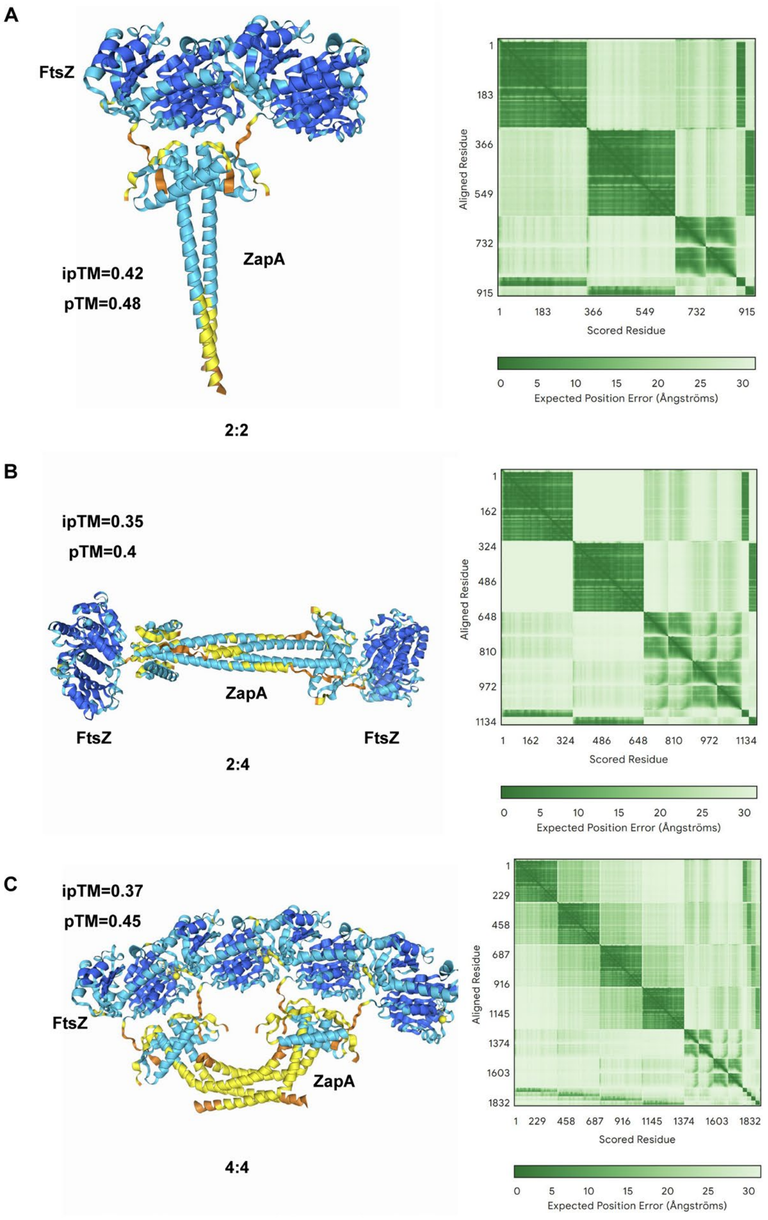


**Fig. S9 Structural models of FtsZ-ZapA complex generated by AlphaFold 3.** (A-C) AlphaFold 3 model of the FtsZ-ZapA complex at ratios of 2:2, 2:4; 4:4 between FtsZ and ZapA. Regardless of the ratio, the N-terminal motif of FtsZ binds to a groove in the dimer head of ZapA tetramer. ipTM, pTM values and PAE for each model were indicated and shown.

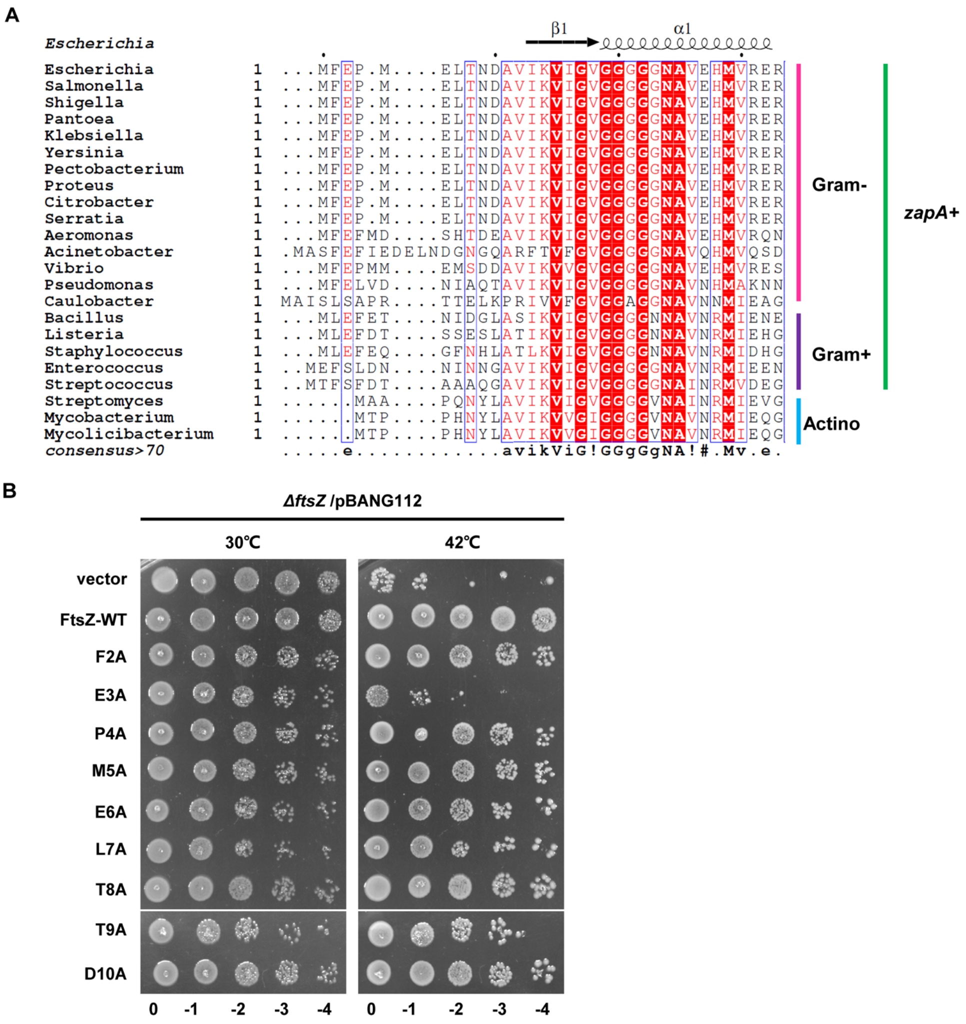


**Fig. S10 Alignment of the N-terminal sequence of FtsZ proteins and complementation test of FtsZ mutants.** (A) Alignment of the N-terminal amino acid sequences of FtsZ proteins from diverse bacteria. In both Gram+ and Gram- bacteria harboring *zapA*, a conserved motif is present at the N-terminus, featured by a large hydrophobic amino acid (F/L) and a glutamate at the second and third position (*E. coli* numbering), respectively, whereas in actinobacteria which do not encode for ZapA these residues are missing. (B) Complementation test of FtsZ N-terminal mutants. Plasmid pKD3C carrying *ftsZ* is temperature sensitive for replication, so it only complements the *ftsZ* defective strain S17 at 30°C. pBANG112 or its derivatives were transformed into S17/pKD3C at 30°C. Transformants were subjected to a spot test on plates at 30 °C or 42°C overnight.

**Table S1. FtsZ mutants isolated by genetic selection**.

| Mutant number | *ftsZ* alleles | Complementation | Resistance to ZapA (IPTG: μM) |
|---|---|---|---|
| M1 | V128I | YES | 1000 |
| M2 | K140E | YES | 125 |
| M3 | G191C,A257T | YES | 1000 |
| M4 | R214H | YES | 125 |
| M5 | T291N | YES | 1000 |
| M6 | T215A,D318N | YES | 125 |
| M7 | N289D | YES | 125 |
| M8 | G94R,A184T,T309I | YES | 125 |
| M9 | K66R,A185V | YES | 1000 |
| M10 | S218Y | YES | 1000 |
| M11 | L248M | YES | 1000 |
| M12 | A184T,V193M | YES | 1000 |
| M13 | V193M | YES | 1000 |
| M14 | K141N | YES | 1000 |
| M15 | L178M,S218F | YES | 1000 |

**Table S2. FtsZ single substitutions with significant resistance to ZapA overexpression.**

| *ftsZ* alleles | Complementation | Resistance to ZapA (IPTG: µM) | Resistance to ZapC | Source |
|---|---|---|---|---|
| WT | YES | NO | NO | Collection |
| V128I | YES | 1000 | NO | This study |
| K141N | YES | 1000 | NO | This study |
| V193M | YES | 1000 | NO | This study |
| S218F/Y | YES | 1000 | NO | This study |
| E219A | YES | 1000 | NO | This study |
| L248M | YES | 1000 | NO | This study |
| A257T | YES | 1000 | NO | This study |
| R258E | YES | 1000 | NO | This study |
| T291N | YES | 1000 | NO | This study |
| K66R | YES | 125 | NO | This study |
| K140E/D | YES | 125 | NO | This study |
| R214H | YES | 125 | NO | This study |
| T215A | YES | 125 | NO | This study |
| N289D | YES | 125 | NO | This study |

**Table S3. List of strains used in this study**.

| Strain | Description | Source/Reference |
|---|---|---|
| BL21 | *F – ompT hsdSB (rB- mB-) gal dcm* (DE3) | Lab collection |
| BTH101 | *F- cya-99, araD139 galE15 galK16 rpsL1 (StrR) hsdR2 mcrA1 mcrB1* | Lab collection |
| CYa35 | W3110, *ftsZ*$^0$, *zipA-mcherry-spc* / pACYC, *ftsZ* & pEXT22, $P_{tac}$*::zapA* | This study |
| CYb1 | W3110,*ftsZ*$^0$ *,zapA-gfp-cat, zapB::kan* /pACYC, *ftsZ* | This study |
| CYm141 | W3110, *ftsZ::cam, zapA<>frt* /pACYC, *ftsZ*$^{N73C}$ | This study |
| CYm142 | W3110/pZH509,*ftsZ-linker5-mNG* | This study |
| CYm143 | W3110/pZH509,*ftsZ-linker5-mNG*&pEXT22,$P_{tac}$*::zapA* | This study |
| EC436 | MC4100, *(λattL-lom)::bla lacIqP*$_{207}$*-gfp-ftsI* | Lab collection |
| JS238 | MC1061, *malPp::lacIq srlC::Tn10 recA1* | Lab collection |
| S17 | W3110, *ftsZ*$^0$ | Lab collection |
| W3110 | *F- λ- rph-1 INV(rrnD, rrnE)* | Lab collection |

**Table S4. List of plasmids used in this study**.

| Plasmid | Description | Source/Reference |
|---|---|---|
| pBANG112 | pACYC, *ftsZ*, $Amp^r$ | (1) |
| pBC1 | pACYC, *ftsZ*$^{N73C}$, $Amp^r$ | This study |
| pCY52 | pE-SUMO,$P_{T7}$*::his-SUMO-ZapA*, $Amp^r$ | This study |
| pCY54 | pE-SUMO,$P_{T7}$*::his-SUMO-ftsZ*$^{L178E}$, $Amp^r$ | This study |
| pCY55 | pE-SUMO,$P_{T7}$*::his-SUMO-ftsZ*$^{D212N}$, $Amp^r$ | This study |
| pCY83 | pKNT25, $P_{lac}$*::ftsZ*$^{1\text{-}316}$*-T25*, $Kan^r$ | This study |
| pCY86 | pKNT25, $P_{lac}$*::ftsZ*$^{317\text{-}383}$*-T25*, $Kan^r$ | This study |
| pCY170 | pEXT22, $P_{tac}$*::zapA-gfp*, $Kan^r$ | This study |
| pCY173 | pEXT22, $P_{tac}$*::zapA*$^{C19A,T50C}$*-gfp*, $Kan^r$ | This study |
| pCY205 | pZH509*, gfp-ftsI,cat,* $Amp^r$ | This study |
| pDSW210 | pDSW206-MCS-*gfp*, $Amp^r$ | (2) |
| pE-SUMO-Amp | pUC, $P_{T7}$*::6XHis-SUMO-MCS*, $Amp^r$ | Lab collection |
| pE-SUMO-FtsZ | pE-SUMO,$P_{T7}$*::his-SUMO-ftsZ*, $Amp^r$ | Lab collection |
| pKD3C | pSC101, *repA*$^{ts}$ *ftsZ* $Cam^r$ | (3) |
| pKNT25 | $P_{lac}$*::T25*, $Kan^r$ | Lab collection |
| pSD319 | pEXT22, $P_{tac}$*::zapA*, $Kan^r$ | Lab collection |
| pSD320 | pEXT22, $P_{tac}$*::zapC*, $Kan^r$ | Lab collection |
| pSY3 | P15A, bla terR $P_{LtetO\text{-}1}$::*ftsZ-mneongreen* | This study |
| pT18A | pUT18, $P_{lac}$*::zapA-T18*, $Amp^r$ | This study |
| pT18-zip | pBluescript II KS, T18-leucine zipper | (4) |
| pT25-zip | pACYC184, *lac UV5T25-leucine zipper* | (4) |
| pUT18C | *Plac::T18*, $Amp^r$ | Lab collection |

| | | |
|---|---|---|
| pXY027 | ColE1, *cat lacI*$^{Q}$ $P_{T5lac}$::*ftsZ-gfp* | (5) |
| pYD208 | P15A, *bla terR* $P_{LtetO-1}$::*ftsW-mneongreen* | Lab collection |
| pZH509-gfp | P15A, *bla terR* $P_{LtetO-1}$*::gfp* | (6) |
| pZT25 | pKNT25, $P_{lac}$*::ftsZ-T25*, Kan$^{r}$ | Lab collection |

**Table S5. Primers used in this study**

| Primer name | Sequence |
|---|---|
| FtsZ-N73C-F | CTGGGCGCTGGCGCTTGTCCAGAAGTTGGCCGC |
| FtsZ-N73C-R | GCGGCCAACTTCTGGACAAGCGCCAGCGCCCAG |
| T18-ZapA-F | CGCGGATCCATCTGCACAACCCGTCGATAT |
| T18-ZapA-R | CGGGGTACCTCATTCAAAGTTTTGGTTAG |
| BsaI-ZapA-F | CGGGTCTCTAGGTTCTGCACAACCCGTCGATAT |
| XbaI-ZapA-R | CGCTCTAGATTATTCAAAGTTTTGGTTAGTTTTTTCGGT |
| FtsZ-L178E-F | GGCCGCGGTATCTCCGAGCTGGATGCGTTTGGC |
| FtsZ-L178E-R | GCCAAACGCATCCAGCTCGGAGATACCGCGGCC |
| FtsZ-D212N-F | GAACGTGGACTTTGCAAACGTACGCACCGTAATG |
| FtsZ-D212N-R | CATTACGGTGCGTACGTTTGCAAAGTCCACGTTC |
| SalI-ZapA-F | ACGCGTCGACGACGTCTCTGCACAACCCGTCGATAT |
| ZapA-HindIII-R | CCCAAGCTTAAAGTGTCATTCAAAGTTTTGGTTAGTTTT |
| pZT25-F | CCCAAGCTTGTAGGCGACAGGCACAAATCGGAG |
| 316-XbaI-R | CGTCTAGAGCGCCGATACCTGTCGCAAC |
| HindIII-317-F | GCAAGCTTATGGGCATGGACAAACGTCCT |

| | |
|---|---|
| 383-XbaI-R | CGTCTAGAGCATCAGCTTGCTTACGCAG |
| XbaI-GFP-F | CGTCTAGAATGAGTAAAGGAGAAGAACTTTTCAC |
| FtsI-HindIII-R | CGAAGCTTGTCGCGCAAATTACGATCTG |
| EcoRI-ZapA-F | CGGAATTCTAATCAATCAGCAGGAAGGT |
| ZapA-GFP-R | CTTCTCCTTTACTCATCGAAAAGTGTTCAAAGTTTTGG |
| ZapA-GFP-F | CTTTGAACACTTTTCGATGAGTAAAGGAGAAGAACTTTTC |
| GFP-HindIII-R | CGAAGCTTTTATTTGTATAGTTCATCCATGCCATG |
| 509-GFPU-F | CATTAAAGAGGAGAAAGGTACCATGAGTAAAGGAGAAGAACT |
| FtsI-C-R | CCAGTGATTTTTTTCTCCATTCTAGATTACGATCTGCCACCT |
| FtsI-C-F | AGGTGGCAGATCGTAATCTAGAATGGAGAAAAAAATCACTGG |
| C-509U-R | TAGCACGCGTACCATGGGATCCTTACGCCCCGCCCTG |
| ZapA-C19A-F | GTTCACTGCGTGTGAACGCGCCGCCTGACCAAAGG |
| ZapA-C19A-R | CCTTTGGTCAGGCGGCGCGTTCACACGCAGTGAAC |
| ZapA-T50C-F | CACTAGAGTCACAAATTGTGAACAGTTGGTCTTC |
| ZapA-T50C-R | GAAGACCAACTGTTCACAATTTGTGACTCTAGTG |
| FtsZ-F2A-F | CAAATCGGAGAGAAACTATGGCGGAACCAATGGAACTTACC |
| FtsZ-F2A-R | GGTAAGTTCCATTGGTTCCGCCATAGTTTCTCTCCGATTTG |
| FtsZ-V128I-F | GATTTGGGTATCCTGACCATTGCTGTCGTCACTAAGC |
| FtsZ-V128I-R | GCTTAGTGACGACAGCAATGGTCAGGATACCCAAATC |
| FtsZ-K141N-F | CAACTTTGAAGGCAAGAATCGTATGGCATTCGCG |
| FtsZ-K141N-R | CGCGAATGCCATACGATTCTTGCCTTCAAAGTTG |
| FtsZ-E219A-F | CGCACCGTAATGTCTGCGATGGGCTACGCAATG |

| | |
|---|---|
| FtsZ-E219A-R | CATTGCGTAGCCCATCGCAGACATTACGGTGCG |
| FtsZ-R258E-F | GACCTGTCTGGCGCGGAAGGCGTGCTGGTTAACATC |
| FtsZ-R258E-R | GATGTTAACCAGCACGCCTTCCGCGCCAGACAGGTC |
| oSY1 | GTGAGCAAAGGCGAAGAAGATAAC |
| oSY2 | GCGGTACCTTTCTCCTCTTTAATG |
| oSY3 | GAGGAGAAAGGTACCGCATGTTTGAACCAATGGAACTTAC |
| oSY4 | TTCGCCTTTGCTCACCCTTCGAATGTGTATCTTGCCTATTCT GCCATCAGCTTGCTTACGC |
